# PKC-dependent enhancement of glutamate input to VTA dopamine neurons in 3xTg-AD mice

**DOI:** 10.1101/2025.07.24.666429

**Authors:** Harris E. Blankenship, Matthew H. Higgs, Kylene M. Harold, Kenneth M. Humphries, Michael J. Beckstead

## Abstract

A growing body of work has recently linked ventral tegmental area (VTA) dopamine neuron dysfunction to Alzheimer’s disease (AD). Work in AD mouse models suggests that VTA dopamine neurons are intrinsically hyperexcitable, yet release less dopamine and exhibit disrupted downstream signaling. Significant focus has been placed on describing dopamine release in projection regions; however, dopamine neurons somatodendritically integrate vast synaptic input, altering action potential output and ultimately determining neurotransmitter release. Synaptic transmission is broadly disrupted in AD, but it is not known to what extent excitatory and inhibitory inputs to the VTA are altered. Here we describe enhanced synaptic excitation in dopamine neurons in the amyloid + tau-driven 3xTg-AD mouse model. Patch-clamp electrophysiology experiments revealed enhanced AMPAR-mediated excitatory input in a subset of perisomatic connections. In contrast, GABA_A_R-mediated inhibition was decreased as a function of dendritic atrophy, determined by single neuron reconstructions. We also detected elevated protein kinase C (PKC) substrate levels in the midbrain, and pharmacological experiments suggested that the strengthened excitation depends on elevated PKC activity. Additionally, postsynaptic AMPA receptor conductance was enhanced and displayed diminished ability to induce plasticity (long-term depression), but this was not dependent upon increased AMPA receptor expression. Morphologically detailed biophysical modeling predicted that synaptic changes, in combination with altered dendritic morphology and intrinsic hypersensitivity, produce increased spontaneous firing rates and a steeper input-output relationship in 3xTg-AD neurons. The results argue against a uniform decrease in synaptic connectivity across the brain in AD, and sheds light on the involvement of deep brain circuits. AD pathology is therefore associated with increased sensitivity of single dopamine neurons, which may help to maintain phasic dopamine signaling in early stages of degeneration.

## Introduction

Growing evidence implicates a variety of cortical and subcortical areas in the pathology and neurodegeneration associated with Alzheimer’s disease (AD). Accumulation of extracellular β-amyloid plaques and intracellular, hyperphosphorylated tau tangles classically define AD pathology(*1, 2*). Clinically, late-stage AD patients present with dementia characterized by memory loss and cognitive decline widely attributed to synaptic dysfunction and neuronal loss in corticolimbic regions(*3, 4*). Research has predominantly focused on the pathological hallmarks of AD in brain regions associated with memory and cognition, notably the hippocampus and cortex. This line of research has advanced our understanding of AD pathophysiology, but has not yielded highly efficacious treatments or a cure for the disease(*5*). Major challenges impeding the development of effective interventions include an incomplete understanding of brain-wide effects and early neurophysiological changes that may precede classical neuropathology.

Recent studies investigating a wide range of brain areas have revealed that cellular findings in hippocampus and cortex are not uniform throughout the brain, and observations in corticolimbic structures cannot be generalized to deeper brain structures(*6–9*). Our group and others have focused on dopamine-releasing neurons in the ventral tegmental area (VTA)(*10–14*), as dysfunction here may help explain common AD comorbidities such as depression and agitation, as well as deficits in cortical and hippocampal plasticity (*11, 12, 15, 16*). These studies implicate dopaminergic alterations early in disease progression, which may contribute to brain-wide pathophysiology(*17–19*).

In rodent models of AD, the autonomous firing of dopamine neurons is elevated due to decreased calcium-activated potassium (SK) channel conductance(*10, 13*). *In vivo*, VTA dopamine neurons receive input through thousands of excitatory, inhibitory, and neuromodulatory synapses that emanate from dozens of brain nuclei(*20, 21*). The currents produced at these synapses integrate with intrinsic conductances to speed or slow dopamine neuron firing, and to drive burst firing(*22–29*), which increases dopamine release locally(*30–33*) and in projection regions(*22, 34–38*). This transient activity of dopamine neurons in response to behaviorally relevant stimuli positions them as a central hub of reward learning(*22*). Recent work in AD models describes mechanisms through which altered VTA dopamine release can adjust learning rules in the hippocampus(*11, 16*). However, to date, no study has investigated how AD pathology affects the synaptic input to VTA dopamine neurons.

Here, we explored synaptic inputs to VTA dopamine neurons in the triple transgenic (3xTg) mouse model of AD, which features both extracellular β-amyloid plaques and intracellular tau tangles in other brain areas(*39*). Using *ex vivo* single-cell patch-clamp electrophysiology, we identified changes in GABAergic and glutamatergic synaptic input, which appear to result from pre- and post-synaptic structural and molecular changes. Notably, dopamine neurons from 3xTg mice exhibited decreased inhibitory input and enhanced excitatory input, both of which contributed to dopamine neuron hyperexcitability. Morphological reconstructions indicated profound dendritic retraction in 3xTg dopamine neurons, suggesting a structural correlate for decreased inhibitory afferent connectivity. We then constructed a family of compartmental biophysical models incorporating altered morphology, previously described intrinsic hypersensitivity(*10*), and newly identified alterations in excitatory and inhibitory input. Simulations predicted that 3xTg dopamine neurons would have an elevated firing rate in response to the same inputs from presynaptic excitatory and inhibitory afferents. In addition, we predict that 3xTg neurons have enhanced sensitivity to presynaptic firing, compared to wild type (WT) neurons. These findings describe enhanced dynamic firing activity in dopamine neurons that may compensate for axonal degeneration in early stages of AD.

## Results

### Reduced VTA dopamine neuron branching in 3xTg mice

In some neurodegenerative models, synapse loss is associated with dendritic hypotrophy(*40–42*). We and others have observed robust dendritic retraction in SNc dopamine neurons in Parkinson’s disease mouse models(*41, 42*), although WT SNc neurons show no change in dendritic structure with healthy aging(*43*). These findings suggest that altered dendritic architecture is driven by pathology and may be a common dopamine neuron response to insult. In a previous study, we also determined that at 12 months of age, VTA dopamine neuron number does not decline in the 3xTg model(*10*). However, immunohistochemistry showed a decrease in tyrosine hydroxylase expression in the region(*10*), which may result from dendritic loss. VTA dopamine neurons typically extend several primary dendrites in the horizontal plane(*44*). To determine if dendrites are altered in brain slices from 3xTg mice, we used a patch clamp electrode to fill individual VTA dopamine neurons with biocytin, then analyzed morphology from skeletonized reconstructions and assessed neurite branching and length (**Figure 1A, Figure S1**). We found that 12 mo 3xTg dopamine neurons exhibit reduced dendritic complexity as assessed by Sholl analysis(*45*) (**Figure 1B-C**), which counts dendritic intersections as a function of distance from the soma(*46*). We also found a significant decrease in total surface area (**Figure 1D**) and cell volume (**Figure 1E**) in 3xTg dopamine neurons compared to WT. Decreased dendritic arborization was previously described in hippocampal pyramidal neurons in an amyloid-only mouse model of AD and resulted in substantial changes in synaptic connectivity and integration(*40*). A diminished somatodendritic surface area could reflect decreased synaptic connectivity and may produce differential effects on synaptic strength.

**Figure 1.**
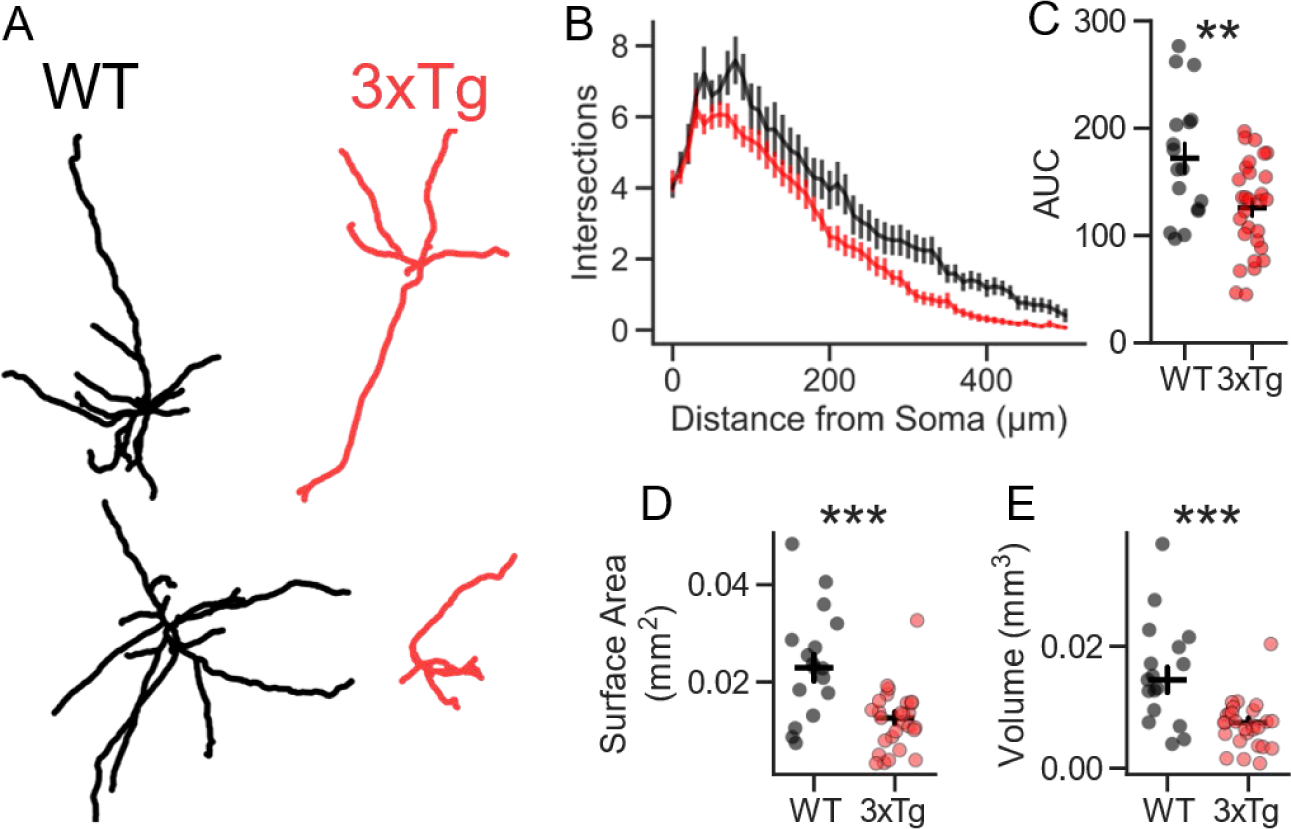
3xTg dopamine neurons exhibit dendritic retraction. **A.** Representative reconstructions of WT (black) and 3xTg (red) dopamine neurons. **B.** Sholl analysis demonstrates decreased dendritic complexity in 12 mo 3xTg dopamine neurons (mixed linear effects regression genotype-radius interaction P=0.022; n=13 [WT] and 17 [3xTg]). **C.** Area under the curve (AUC) for the Sholl plots was lower in 3xTg neurons (two-tailed T-test, t=3.08, P=0.0036). **D.** Total surface area was lower in 3xTg neurons (Mann-Whitney, U=374.0, P=0.00052). **E.** Total cell volume was lower in 3xTg neurons (Mann-Whitney, U=378.0, P=0.00036).

### Enhanced synaptic excitation in 3xTg dopamine neurons

In AD models, synaptic input to neurons might be altered by signaling cascades triggered by amyloid and/or tau, or as a homeostatic adaptation to changes in neuronal activity(*47–50*). The latter possibility, coupled with decreased cellular surface area, suggests that VTA dopamine neurons may exhibit dampened excitatory synaptic input in the 3xTg model as a homeostatic response to their intrinsic hyperexcitability(*10*). To assess the overall change in excitatory input to these cells, we recorded AMPA receptor (AMPAR)-mediated miniature excitatory postsynaptic currents (mEPSCs) isolated in the presence of the sodium channel blocker tetrodotoxin (TTX) and neurotransmitter receptor blockers picrotoxin (GABA_A_R), MK-801 (NMDAR), hexamethonium (nicotinic acetylcholine receptors), and CGP55845 (GABA_B_R). We recorded from 3 mo and 12 mo 3xTg and non-transgenic (WT) VTA dopamine neurons which coincided prior to and concurrent with our previous observations of hyperexcitability, respectively. Surprisingly, we found that mEPSC amplitudes were elevated in both 3 mo and 12 mo 3xTg mice (**Figure 2A-B**), in contrast to our homeostatic plasticity hypothesis. Furthermore, mEPSC frequency was enhanced in 12 mo 3xTg dopamine neurons (**Figure 2C**). To ensure that this finding was not merely due to an inability to identify small synaptic events, we downscaled mEPSCs from WT mice, added a noise trace obtained in the presence of the AMPAR blocker DNQX, and re-ran them through our detection algorithm (**Figure S2A-F**). While the algorithm successfully detected the decrease in amplitude (**Figure S2G**) there was no detectable difference in the number of events detected (**Figure S2H**), indicating that observed differences in mEPSC frequency were biological. The sustained, progressive changes in synaptic excitation observed in 3xTg dopamine neurons differ from recent findings in hippocampal pyramidal neurons in the amyloid-mutation knock-in APP^NL-G-F^ mice, which indicate homeostatic flexibility in excitatory transmission as disease progresses(*6*).

**Figure 2.**
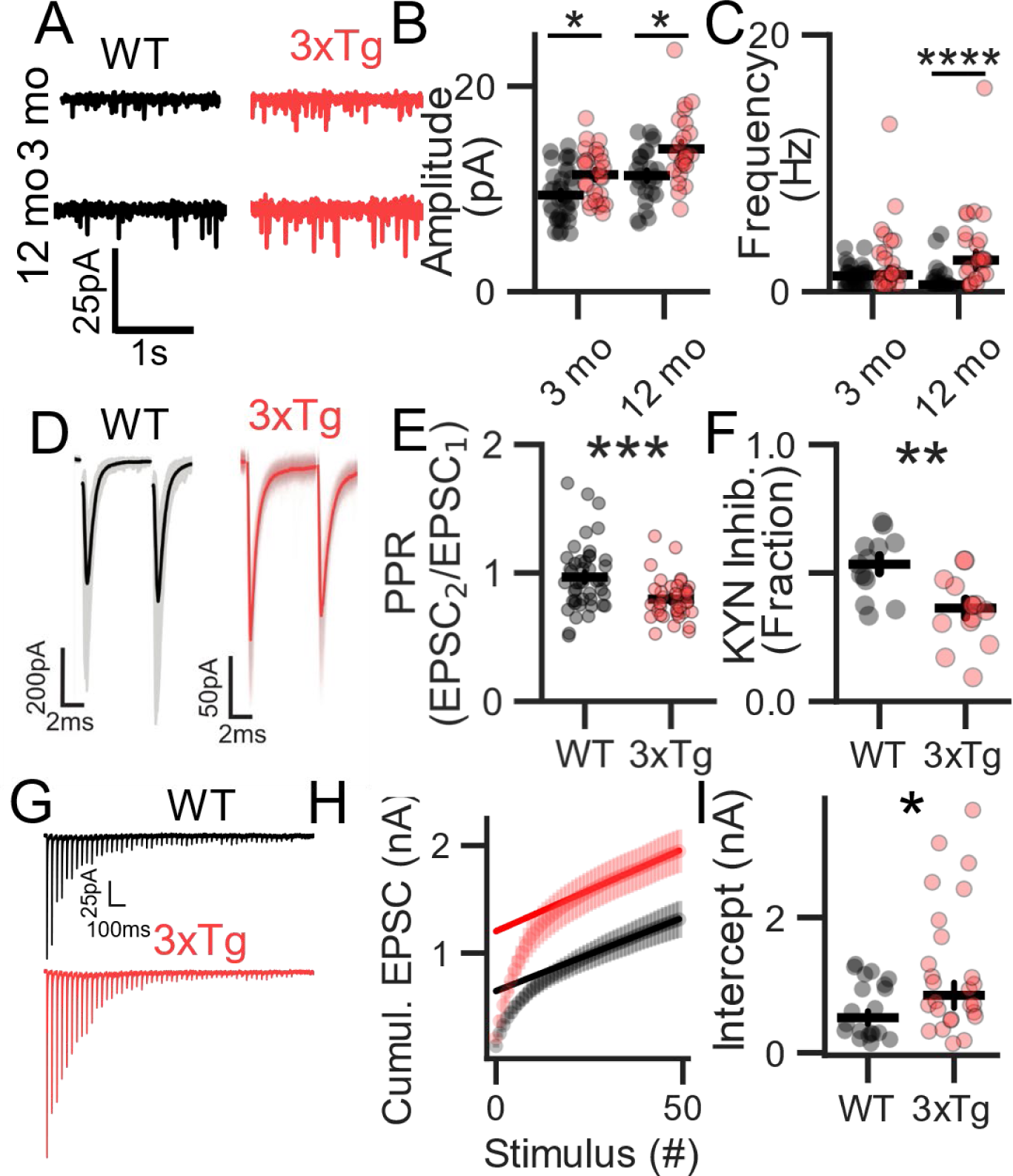
Enhanced excitatory synaptic input in 3×Tg VTA dopamine neurons. **A.** Representative traces of mEPSCs in 3 and 12 mo WT and 3xTg neurons. **B.** mEPSC amplitude is increased in 3xTg mice at 3 months (Mann-Whitney, Sidak post hoc P=0.0109; n=35 [WT] and 26 [3xTg]) and 12 months (Mann-Whitney, Sidak post hoc, P=0.016; n=26 [WT] and 24 [3xTg]) (2-way ANOVA, main effects of age [P=6.484×10^-5^, F=17.29] and genotype [P=3.32×10^-5^, F=18.85]). **C.** Increased mEPSC frequency in 12 mo 3xTg neurons (Mann-Whitney, Sidak post hoc P=2.85 x10^-5^, 2-way ANOVA, main effect of genotype P=3.82×10^-5^, F=18.47). **D-E.** Smaller paired-pulse ratios of evoked EPSCs in 12 mo 3xTg dopamine neurons (Mann-Whitney, U=1266.0, P=0.005; n=45 [WT] and 39 [3xTg]). **F**. Diminished inhibition of EPSC amplitude by kynurenic acid (250 μM) in 3xTg mice (two-tailed t-test, t=-3.56, P=0.0015; n=15 [WT] and 14 [3xTg]). **G.** Representative EPSC trains evoked by 20 Hz stimuli in a WT neuron (top) and a 3xTg neuron (bottom). **H-I.** Enlarged readily releasable pool, assayed by the asymptotic y-intercept of the cumulative EPSC amplitude plot (Mann-Whitney, U=170.0, P=0.0473; n=20 [WT] and 26 [3xTg]).

The enhancement of mEPSC frequency in 12 mo 3xTg mice may suggest a presynaptic locus. The corresponding effect on evoked release might then be an increase in the release probability of each synaptic vesicle and/or an increase in the readily releasable pool (RRP) of vesicles. A recent report indicated that β-amyloid oligomers increase the probability of release at glutamatergic synapses(*51*). To assess this possibility in VTA dopamine neurons, we evoked EPSCs with a bipolar stimulating electrode at 20 Hz in slices from 12 mo WT and 3xTg mice, and calculated the paired pulse ratio (PPR), a measure inversely correlated to initial release probability(*52*). The PPR was lower in 3xTg dopamine neurons, consistent with a higher initial release probability (**Figure 2D,E**)(*52*).

If the probability of vesicle release at individual synapses is increased, this might result in multivesicular release and consequent elevation of synaptic glutamate concentrations. At other central synapses, increasing the probability of release and the RRP elevates synaptic glutamate concentrations, which can be measured using AMPA receptor competitive antagonism(*53–55*). To determine if glutamatergic afferents in 3xTg mice generate elevated synaptic glutamate concentrations, we evoked EPSCs and washed on kynurenic acid (200 μM), a competitive antagonist of AMPA receptors. Kynurenic acid blocked a smaller fraction of the EPSCs in the 3xTg neurons than in WT neurons, consistent with enhanced glutamate release (**Figure 2F**).

To determine if the RRP in the stimulated afferents is altered in 3xTg mice, we delivered stimulus trains at 20 Hz to deplete the pool (**Figure 2G**). To estimate the RRP size, we plotted the cumulative EPSC as a function of stimulus number, obtained a linear fit to the late portion of the plot, and measured the RRP as the y-intercept of the fit line (**Figure 2H**)(*56–58*). This analysis demonstrated that the RRP was larger in 12 mo 3xTg dopamine neurons compared to WT (**Figure 2I**). The apparent change in the RRP, quantified as total current amplitude, was larger than the enhancement of mEPSC amplitude described above, suggesting that the number of synaptic vesicles making up the RRP was increased. These data suggest that in 3xTg neurons, the set of glutamatergic afferents activated at a given stimulus intensity contained more release-competent vesicles at their synapses onto each recorded neuron.

One possible explanation for the elevated pool size is altered presynaptic kinase activity(*54, 59–61*). Synaptic kinase activity is associated with pathological progression in AD(*62–64*), and β-amyloid has been linked to increased pool size and spontaneous synaptic vesicle exocytosis(*65*) as well as elevated presynaptic protein kinase C (PKC)(*51*). Manipulation of presynaptic protein kinase A would be expected to increase the RRP without changing release probability, which did not appear consistent with our findings(*54*). In contrast, phosphorylation of presynaptic proteins by PKC increases release probability(*61, 66, 67*) while enlarging the RRP(*59, 61*) and elevating mEPSC frequency(*61, 68*), which is consistent with our data. Importantly, hyperactivity of PKCα (a member of the diacylglycerol (DAG)-dependent, or ‘conventional’ class) has been detected in AD patients(*62, 63*), and appears to drive cognitive deficits in mouse models(*64*). To determine if PKC expression or activity was increased in 3xTg mice, we carefully isolated tissue from the ventral midbrain (**Figure S3A,B**) and immunoblotted against PKCα and PKC substrates. While there was no significant difference in PKCα (**Figure S3C,D**) or phosphorylated PKC (**Figure S3E,F**), levels of phosphorylated PKC substrates (**Figure 3A,B**) and the ratio of phosphorylated substrates to PKCα (**Figure S3G**) were significantly elevated in the 3xTg ventral midbrain compared to controls.

**Figure 3.**
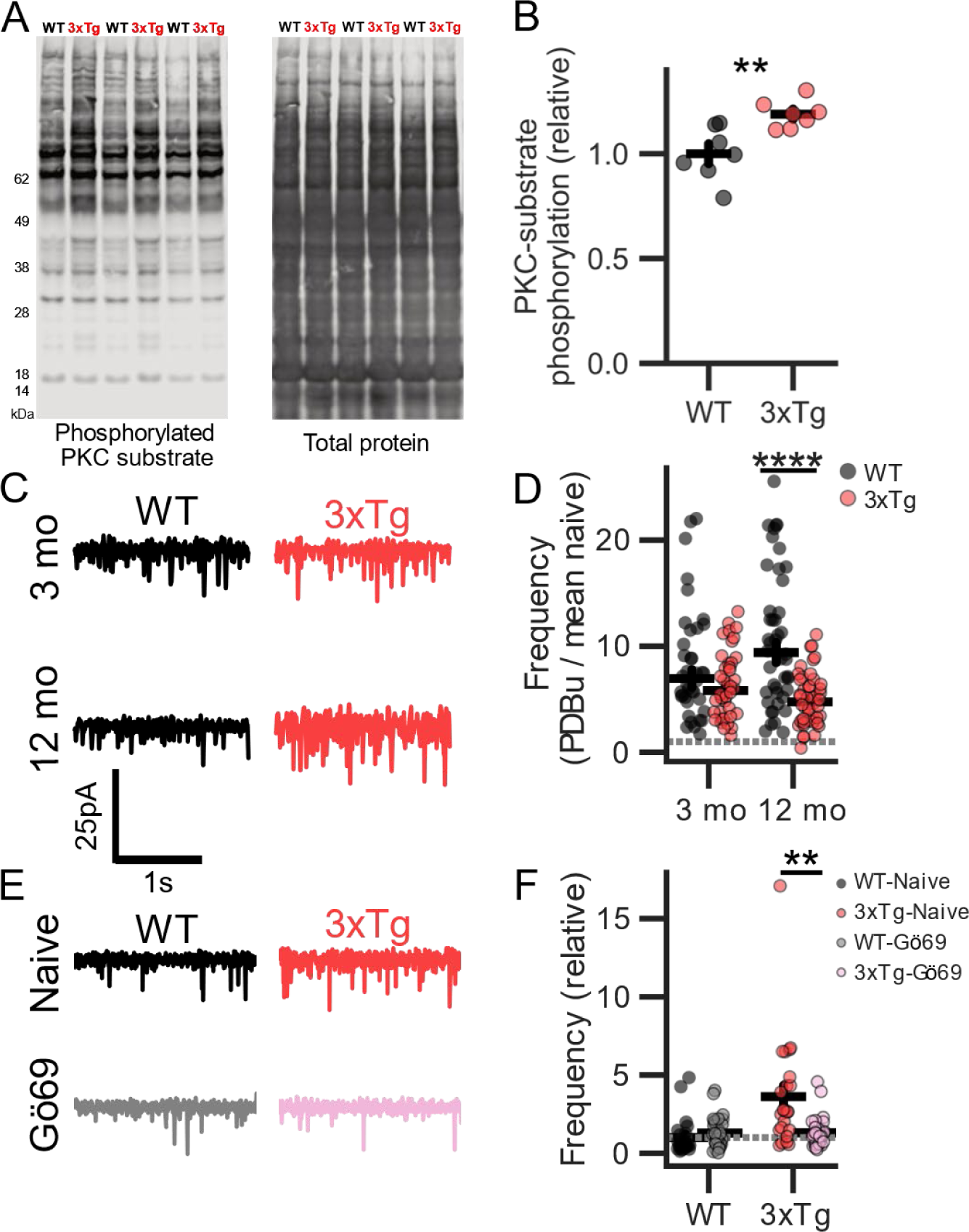
PKC activity drives presynaptic enhancement in 3xTg dopamine neurons. **A.** Western blot of ventral midbrain lysates immunoblotted with a pan-specific antibody against phosphorylated substrates of PKC and stained for lysate total protein. **B**. Quantification of (A). Phosphorylation of PKC substrates is significantly enhanced in the 3xTg ventral midbrain (two-tailed t-test, t=-3.5, P=0.0045; n=14 mice, 7 replicates). **C.** mEPSCs recorded after treatment with PDBu. **D**. mEPSC frequency after PDBu treatment, expressed as frequency relative to the mean value for untreated neurons of the same age and genotype. The relative change by PDBu was smaller in 12 mo 3xTg dopamine neurons (Sidak post-hoc P=7.65×10^-6^, 2-way ANOVA main effect of genotype F=30.64, P=1.21×10^-7^ and genotype-age interaction F=7.06, P=8.63×10^-3^; 3 mo: n=38 [WT] and 40 [3xTg]; 12 mo: n=44 [WT] and 46 [3xTg]). **E.** mEPSCs recorded from naïve and PKC inhibitor (Gö6983) treated 12 mo WT and 3xTg dopamine neurons. **F.** mEPSC frequency expressed as frequency relative to WT-naïve (2-way ANOVA, main effect of genotype, F=11.34, P=0.001; treatment, F=7.59, P=0.0068; and genotype-treatment interaction, F=14.74, P=0.0002; n=37 [WT] and 31 [3xTg]). 12 mo WT dopamine neurons display no effect of PKC inhibition (Sidak’s post-hoc, P=0.144), but a decrease in frequency in the 3xTg group (Sidak’s post-hoc, P=0.008).

To further investigate PKC influence on excitatory input to 3xTg dopamine neurons, we next used a phorbol ester (DAG analog) to target C1 domains on PKC(*69*) and probed the consequences of PKC activation(*59, 70*). We preincubated brain slices with a saturating concentration of the phorbol ester phorbol-12,13-dibutyrate (PDBu, 1 µM)(*71*), and measured mEPSC frequency (**Figure 3C**). In the presence of PDBu, mEPSC frequency was dramatically increased in both WT (**Figure S4A**) and 3xTg (**Figure S4B**) neurons from both age groups. However, the relative enhancement was smaller in the 12 mo 3xTg dopamine neurons (**Figure 3D**). These data suggest that the effect of PDBu on mEPSC frequency was partially occluded in the 3xTg mice, possibly suggesting presynaptic phosphorylation of PKC targets in 3xTg mice(*59, 67, 72, 73*). To determine if inhibition of PKC inhibition could recover mEPSC amplitude in 12 mo 3xTg neurons, we preincubated slices in the PKC specific inhibitor Gö6983 (1 µM) for at least 1 hour prior to recording (**Figure 3E**). PKC inhibition had no significant effect on WT mEPSC frequency, but reduced 3xTg mEPSC frequency to WT levels (**Figure 3F**). Together, these data support a model of elevated presynaptic PKC activity in 3xTg mice resulting in enhanced glutamate release.

### Selective enhancement of AMPAR versus NMDAR EPSCs

The increase in mEPSC amplitude suggests that the postsynaptic AMPA receptor conductance is greater in 3xTg mice(*74, 75*). Thus, if the NMDA receptor conductance is unaffected, we would expect to see an increase in the AMPAR:NMDAR EPSC ratio. To measure this, we voltage clamped cells at +40mV and evoked an outward EPSC comprised of both AMPAR and NMDAR components (**Figure 4A**). After obtaining a stable baseline, we applied the NMDAR-specific antagonist APV (50 µM), leaving only the AMPAR-mediated component. The AMPAR EPSC was subtracted from the dual-component EPSC to obtain the NMDAR EPSC. Dividing the AMPAR EPSC peak amplitude by the NMDAR EPSC peak amplitude revealed a significant enhancement of the AMPAR:NMDAR ratio in 12 mo 3xTg dopamine neurons (**Figure 4B**).

**Figure 4.**
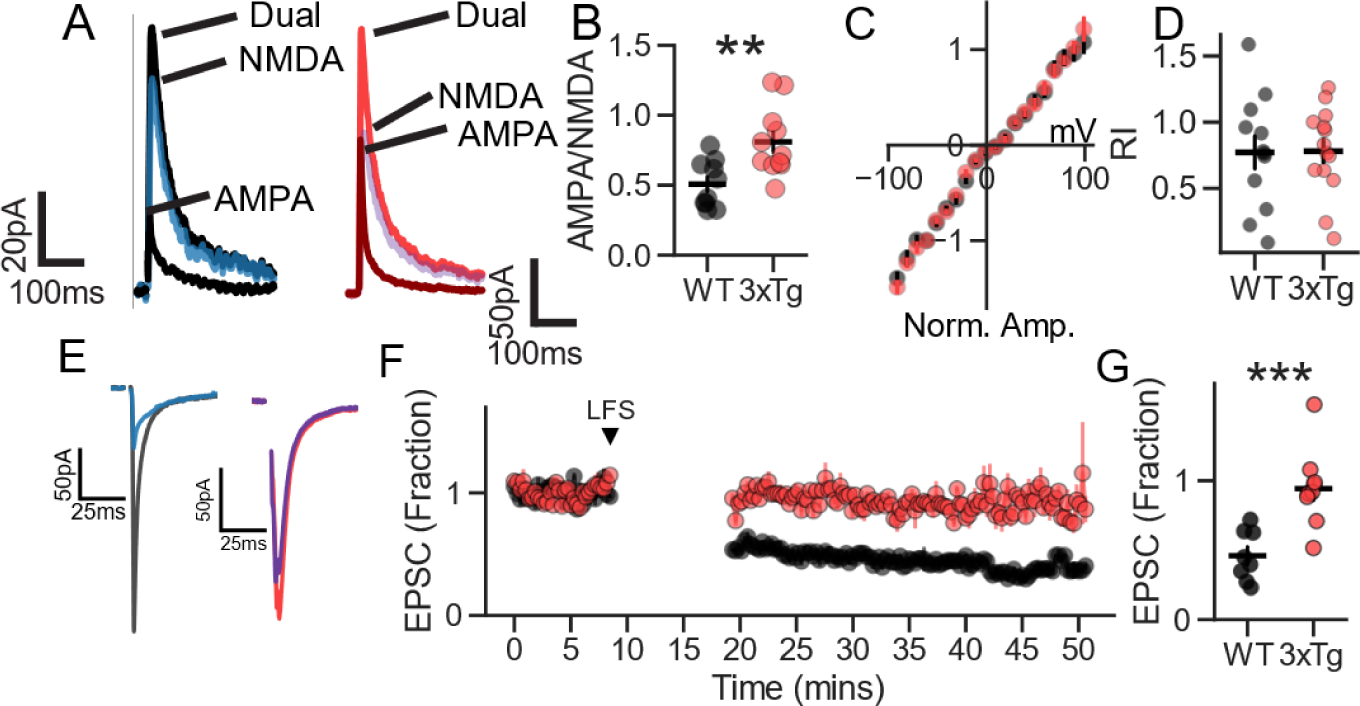
AMPA receptor current enhancement and decreased plasticity in 3xTg dopamine neurons. **A.** AMPAR and NMDAR components of EPSCs evoked at +40 mV**. B.** 12 mo 3xTg dopamine neurons display elevated AMPAR:NMDAR ratios (t=-3.29, P=0.0039; n=10 [WT] and 11 [3xTg]). **C.** Average current-voltage curves for evoked AMPAR EPSCs in WT and 3xTg dopamine neurons, corrected for liquid junction potential. **D.** The rectification index (RI) did not differ between groups (n=12 [WT] and 14 [3xTg]). **E.** Representative EPSCs in WT (black and blue) and 3xTg (red and purple) before and after low frequency stimulation (1Hz for five minutes). **F.** Time course of WT (black) and 3xTg (red) responses before and after LFS. **G.** Quantification of **F.**, showing a significant difference in LTD between the two groups (t=-4.39, P=0.00045; n=11/group).

To test whether EPSC enhancement was due to increased expression of AMPA receptors, but not NMDA receptors, we next measured the response to selective glutamate receptor agonists(*76, 77*). To assay the AMPAR current, we washed on the non-desensitizing AMPA receptor agonist kainic acid (250 µM)(*78*) in the presence of TTX and synaptic blockers. The kainic acid-induced inward current was larger in the 3xTg neurons compared to WT (**Figure S5A,B**), consistent with increased AMPA receptor cell surface expression. In contrast, the current measured in response to the agonist NMDA (50 μM) while holding the neuron at +40 mV did not differ between dopamine neurons from WT and 3xTg mice (**Figure S5C,D**).

Some forms of synaptic potentiation that increase the AMPAR:NMDAR EPSC ratio in VTA dopamine neurons rely on addition of GluA2-lacking, Ca2+ permeable AMPA receptors(*79, 80*) that exhibit inward rectification(*81*). To determine if 3xTg dopamine neurons display AMPA receptor rectification, we calculated current-voltage (I-V) curves for evoked AMPAR-mediated EPSCs in 12 mo WT and 3xTg dopamine neurons (**Figure 4C**). There was no difference in rectification between the two genotypes (**Figure 4D**), suggesting that 3xTg dopamine neurons do not have increased functional expression of GluA2-lacking AMPA receptors.

Treatments that induce AMPAR EPSC potentiation in dopamine neurons, either *in vivo* or *ex vivo*, can later enhance long-term synaptic depression (LTD)(*77, 79*). To determine whether this holds in the 3xTg model, we applied low-frequency stimulation (LFS; 1Hz). Unexpectedly, WT dopamine neurons from 12 mo mice underwent LTD, but 3xTg neurons did not (**Figure 4E-G**), which is opposite of the previously reported effects. These data suggest that the enhancement of AMPAR EPSCs found in 3xTg dopamine neurons differs from previously reported changes produced by *in vivo* cocaine exposure, food restriction, fear learning, and *ex vivo* LTP induction. Further, the EPSC enhancement in the 3xTg model may be relatively resistant to reversal.

### Changes in unitary EPSCs

In most experiments using strong extracellular stimuli, evoked synaptic currents are produced by activation of more than one axon synapsing on the recorded neuron. Comparison of synaptic current amplitudes then must assume that equivalent stimuli activate the same fraction of the afferents contacting each neuron. To address this limitation, we next employed minimal electrical stimulation to evoke AMPAR-mediated unitary (u)EPSCs in 12 mo WT and 3xTg dopamine neurons (**Figure 5**). Given the changes in mEPSCs and compound evoked EPSCs described above, we expected to see a general enhancement of uEPSC amplitudes. In the WT group, uEPSC amplitudes had a relatively narrow distribution (**Figure 5A**, mean 27 pA ± 24; SD), whereas the 3xTg uEPSCs displayed a much wider distribution (**Figure 5B**, mean 59 pA ± 82; SD). Interestingly, the difference between genotypes was only evident at large amplitudes, indicated by a striking rightward shift at the top quartile of the cumulative distribution (**Figure 5C**). Plotting uEPSC rise time constants against the corresponding amplitudes, it was apparent that the currents with the largest amplitudes had short rise times (**Figure 5D**). Additionally, while 3xTg uEPSCs had longer decay time constants on average (**Figure S6A**) the largest uEPSCs also had the fastest decay time constants (**Figure S6B**) and the lowest coefficient of variation (CV; **Figure S6C**). This suggests that a majority of uEPSCs, and thus single-axon connections, remain unchanged, but a subpopulation of excitatory inputs are strengthened in 3xTg mice. These inputs most likely arrive near the soma, as their large amplitudes and fast rise time constants suggest minimal electrotonic filtering(*82*).

**Figure 5.**
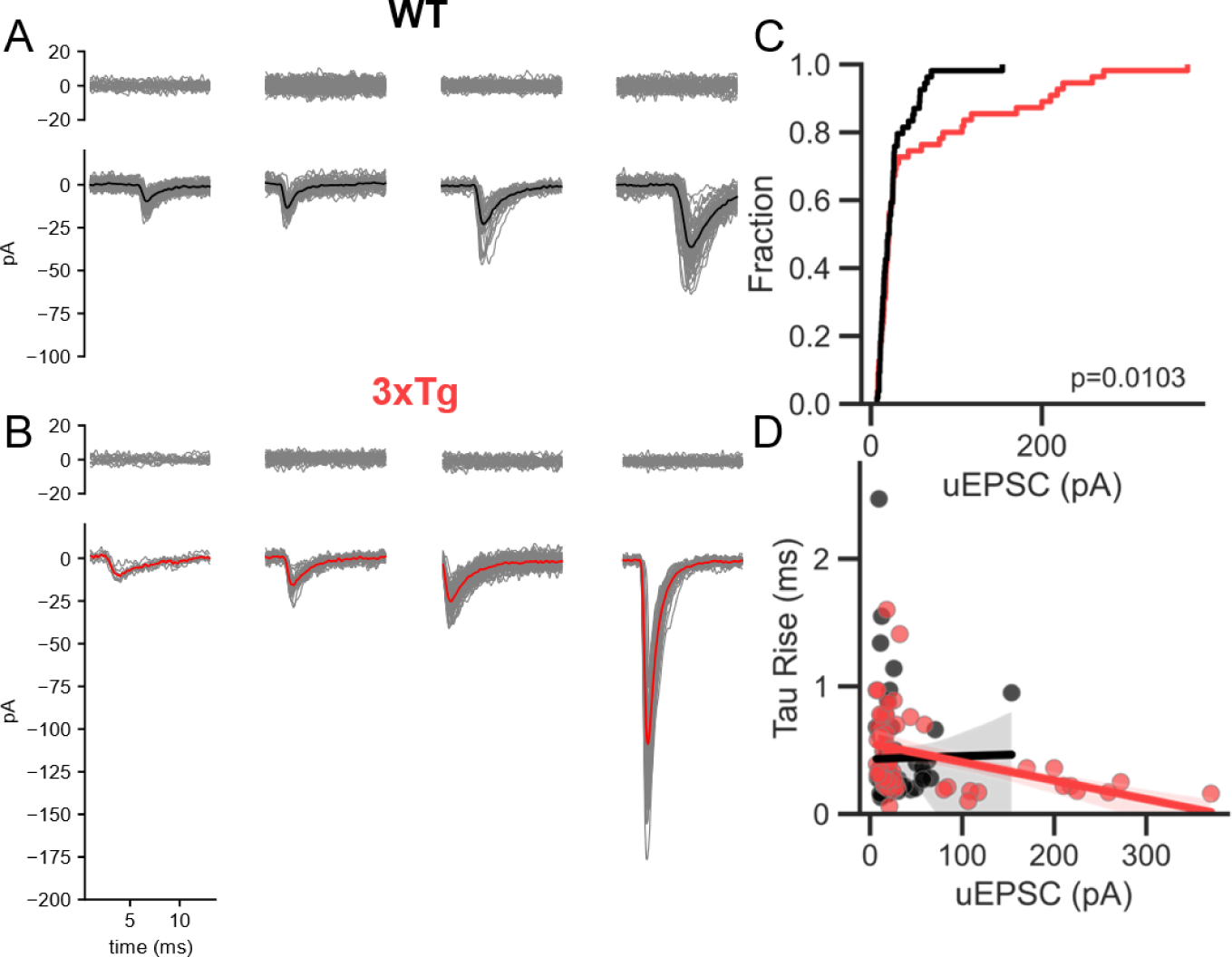
The top quartile of uEPSCs are enhanced in 3xTg dopamine neurons. **A.** Representative uEPSCs from WT neurons for each quartile of the amplitude distribution (n=57). **B.** Same as A, but for 3xTg neurons (n=71). **C.** Cumulative probability of uEPSC amplitudes for WT (black) and 3xTg (red) neurons. The distributions were significantly different (Singleton-Epps two-sample test, P=0.0103) and showed a clear divergence in the top quartile. **D.** uEPSC rise time constants plotted against the corresponding amplitudes. In the 3xTg group, the large events had fast rise times.

### AMPA receptor expression does not explain heightened synaptic strength

Given the evidence suggesting AMPAR enhancement may be limited to a subset of synapses localized near the soma, we sought to determine the distribution of AMPA receptor expression in WT and 3xTg dopamine neurons. We filled individual neurons with biocytin through a recording pipette, reconstructed their somatodendtic morphology ofline, and performed immunofluorescence against the AMPA receptor 1 (GluA1) subunit(*83*). Twelve-month-old WT and 3xTg dopamine neurons robustly expressed GluA1 across their entire somatodendritic extent (**Figure S7A,B**), consistent with previous reports(*83*). On average, 3xTg dopamine neurons displayed roughly half as many GluA1 puncta as WT (**Figure S7C**; WT: 827 ± 94 puncta/neuron, 3xTg: 434 ± 46 puncta/neuron). Total dendritic length was lower in 3xTg dopamine neurons than in WT on average (**Figure S7D**). The decrease in GluA1 puncta was larger than the reduction in dendritic length, as the density (puncta/µm) of GluA1 puncta was lower in 3xTg dopamine neurons than in WT (**Figure S7E**), but density calculated as a function of surface area was nearly identical (**Figure S8**). Additionally, the average intensity of individual puncta was similar between groups (**Figure S7F**).

To determine whether 3xTg neurons display a shift in GluA1 localization, we skeletonized reconstructions and computed a graph to asses the relationship between distance from the soma and GluA1 intenties (**Figure S7G,H**, see Methods). The relationship between GluA1 intensity in puncta and distance from the soma did not differ between WT and 3xTg neurons (**Figure S7I**), as quantified by the slope of the relationship for each cell (**Figure S7J**). Thus the increased AMPA receptor currents in 3xTg mice cannot be explained by enhanced GluA1 expression.

### 3xTg VTA dopamine neurons lose inhibitory synaptic connectivity

Given the increase in excitatory input, we hypothesized that VTA dopamine neurons in the 3xTg mice might maintain homeostatic regulation of their activity by strengthening inhibitory connections. To assess the overall change in synaptic inhibition, we measured GABA_A_R miniature inhibitory postsynaptic currents (mIPSCs) isolated in the presence of TTX, DNQX, MK-801, hexamethonium, and CGP55845; the recorded currents were completely blocked by the GABA_A_ receptor antagonist picrotoxin (**Figure S9A,B**). The amplitude of mIPSCs did not differ between WT and 3xTg dopamine neurons at either 3 or 12 months of age (**Figure 6A,B**). However, mIPSC frequency was reduced in 3- and 12-mo 3xTg neurons (**Figure 6C**). This might reflect presynaptic changes at individual inhibitory synapses and/or a smaller number of synapses associated with the reduced dendritic arbor. To assess the possibility of a change in release probability, we electrically evoked IPSCs (eIPSCs, **Figure S9C,D**) at 20 Hz and measured PPRs. In 12 mo mice, the age at which the relative difference in mIPSC frequency was largest (**Figure 6B**), we detected no difference in PPR (**Figure 6D,E**), providing no evidence for a change in release probability. The eIPSC amplitudes were lower in 3xTg neurons compared to WT with the same stimulus strength (**Figure 6F**), consistent with less total inhibitory input. The eIPSC decay time constant did not differ between groups (**Figure 6G**).

**Figure 6.**
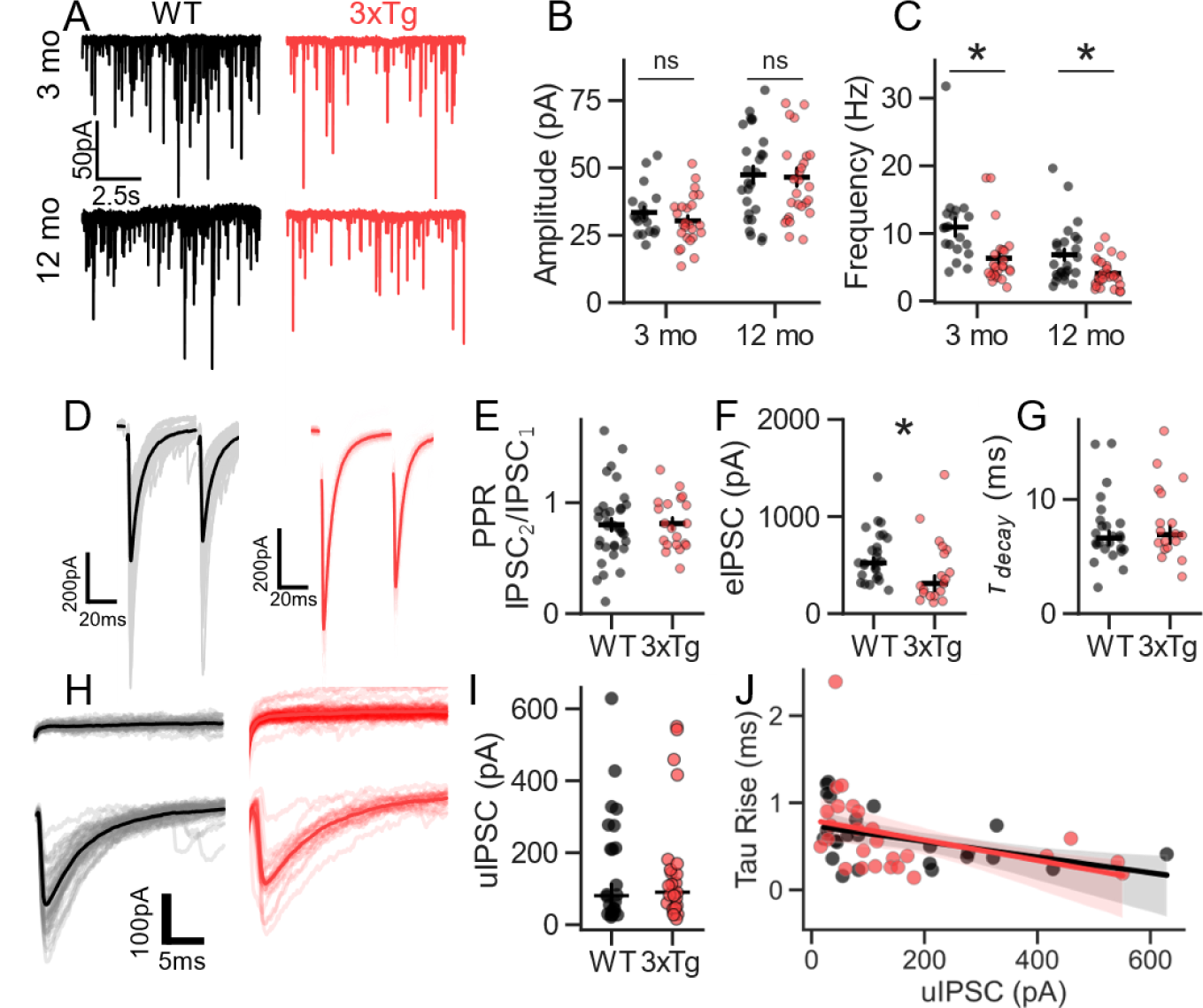
Decreased inhibitory input to 3xTg dopamine neurons. **A.** Representative traces of mIPSCs in 3- and 12-mo WT and 3xTg neurons. **B.** No effect of genotype in mIPSC amplitudes in 3 or 12 mo 3xTg mice (3 mo: n=19 [WT] and 25 [3xTg]; 12 mo: n=26 [WT] and 27 [3xTg]). **C.** Decrease in mIPSC frequency (2-way ANOVA, main effects of age (F=13.1211, P=0.0005) and genotype (F=17.735, P=0.001)) in 3 mo (Sidak post hoc, P=0.014) and 12 mo 3xTg mice (Sidak post hoc, P=0.01). **D-E.** No effect of genotype on 20 Hz paired-pulse ratio. **F.** Smaller average evoked IPSC amplitudes in 3xTg mice (Mann-Whitney, U=148.0, P=0.0236; n=26 [WT] and 19 [3xTg]). **G.** No difference in the decay time constant of evoked IPSCs. **H.** Examples of presumptive unitary IPSCs evoked by minimal electrical stimulation in a WT neuron (left) and a 3xTg neuron (right). Top traces are failures, and bottom traces are successes. Light traces are from individual trials. **I.** uIPSC amplitudes did not differ between WT and 3xTg dopamine neurons (n=24 [WT] and 26 [3xTg]). **J.** Similar negative correlations between rise-time constants and uIPSC amplitudes between WT and 3xTg dopamine neurons.

To determine whether single-axon inhibitory connections are weaker in 3xTg neurons, we employed minimal stimulation to isolate uIPSCs(*84*). In contrast to the results obtained with strong stimuli, we found that uIPSC amplitude (**Figure 6H,I**) did not differ between WT and 3xTg dopamine neurons. Both groups showed similar, skewed distributions of uIPSC amplitudes in line with our previous report(*84*). The relationship between uIPSC amplitudes and rise times was similar between genotypes (**Figure 6J**), providing no evidence for a change in the electrotonic distance of inhibitory synapses from the soma. Taken together, these results suggest that the total number of inhibitory afferents contacting each dopamine neuron is lower in the 3xTg mice, but the strength of the remaining inhibitory synapses and the number formed by each afferent axon are most likely maintained.

### Computational modeling of 3xTg dopamine neuron firing *in vivo*

To predict how the altered synaptic inputs, intrinsic firing properties, and morphology of 3xTg dopamine neurons affect their firing activity *in vivo*, we turned to biophysical modeling. Biophysical models of dopamine neurons can recapitulate morphology, synaptic weights and locations, intrinsic conductances, and passive properties to predict physiologically relevant outcomes(*85–89*). Previously, we reported that dopamine neurons from 3xTg mice have decreased small-conductance calcium-activated potassium channel (SK) calcium sensitivity, leading to increased firing rates and a steeper input-output relationship in response to somatic current injection(*10*). The present study described enhancement of excitatory synaptic weights, reduction of inhibitory afferent connectivity, and morphological retraction in 3xTg mice. We combined these properties to create a family of WT and 3xTg dopamine neuron multi-compartment biophysical models and simulated their firing activity in response to a range of synaptic inputs. The models were constructed using simplified dendritic morphologies derived from 17 reconstructed WT dopamine neurons and 28 3xTg neurons (**Figure S1**) and included ionic conductances known to govern spontaneous firing, including a non-inactivating NALCN-like Na+ conductance, a slowly inactivating A-type potassium conductance, and an SK conductance(*10, 86, 90, 91*)(**Figure S10**). To incorporate our previous findings, we lowered the SK channel conductance in the 3xTg model, decreasing the medium after-hyperpolarization(*10*).

We next performed simulations to investigate the influence of differential synaptic input. The input to each model neuron was simulated by constructing a network of independently firing presynaptic excitatory and inhibitory neurons. The number of neurons in each cell’s presynaptic network was scaled in proportion to the postsynaptic cell’s surface area, where the average WT surface area resulted in 250 presynaptic excitatory afferents and the same number of inhibitory afferents (**Figure 7A,B**). Each presynaptic neuron was stimulated with brief current pulses delivered as a homogenous Poisson point process, each pulse producing one spike (**Figure S11**). It is recognized that many of the single-axon connections producing unitary synaptic currents likely form more than one synapse. However, as the morphological details of these connections are not known, the model configuration was simplified to deliver each unitary synaptic conductance at one site.

**Figure 7.**
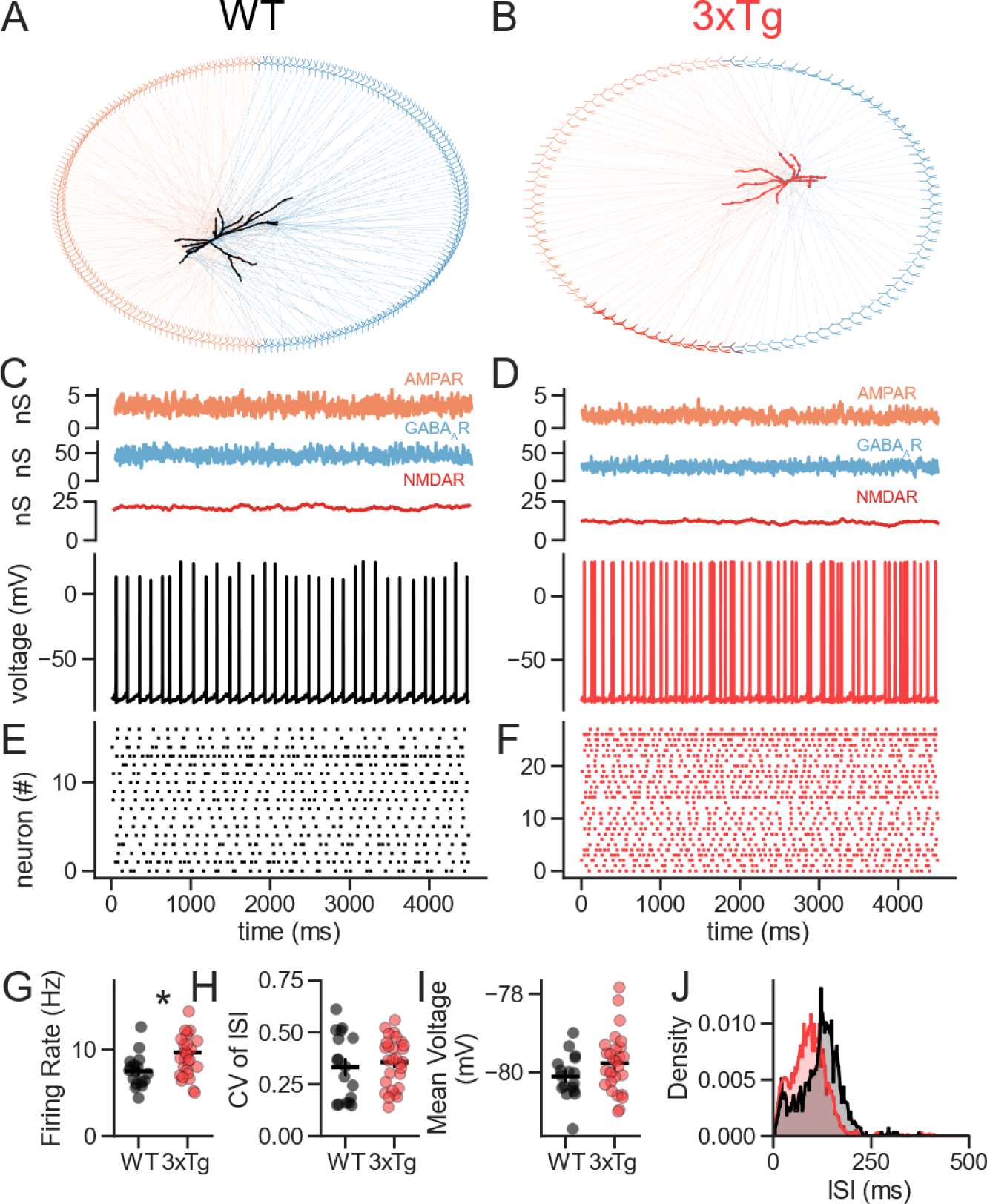
Biophysical modeling predicts that, given the same excitatory and inhibitory afferent firing rates, 3xTg dopamine neurons would fire faster than WT neurons. **A**. Model WT dopamine neuron (black) connected to a presynaptic network of excitatory neurons (orange) and inhibitory neurons (blue). For clarity, the plot illustrates only 100 presynaptic neurons (50 excitatory and 50 inhibitory). **B**. 3xTg network similar to represented with half as many presynaptic neurons A. The subset of excitatory neurons in dark red represents those producing enhanced AMPAR conductance. **C**. Simulation of response to a steady-state barrage of synaptic excitation and inhibition in a WT neuron. Top traces are the summed synaptic conductances for AMPAR (orange), GABA_A_R (blue), and NMDAR (red). Bottom trace is the somatic membrane potential response (black). **D**. Same as C, but for a 3xTg neuron. **E**. Raster plot of all WT spike times in multiple trials of the simulation. **F**. Same as E, but for the 3xTg neuron. **G**. 3xTg compartmental biophysical models display elevated firing rates compared to WT (Mann-Whitney, U=129.0, P=0.011; n=13 [WT] and 17 [3xTg]). **H**. There was no effect of genotype on the CV of the ISI (Mann-Whitney, U=214.0, P=0.58). **I**. There was no difference in the mean somatic membrane potential (t=-1.58, P=0.12). **J**. Histogram of ISIs displayed a leftward shift in the 3xTg group.

To simulate the enhanced synaptic excitation observed in the 3xTg neurons, we quadrupled the AMPAR conductance for the excitatory connections on the soma and on first-order dendrites. With the baseline input described above, the WT dopamine neuron models fired an average of 7.50 spikes/s (±0.47; SEM), and the 3xTg neuron models fired 9.71 spikes/s (±0.65; SEM) (**Figure 7-G**). The regularity of firing was similar between WT and 3xTg models, as measured by the CV of the ISI (**Figure 7H**). Previously, we reported a depolarization of the ISI voltage during spontaneous firing of 3xTg neurons(*10*); however, the average membrane potential was not significantly different between modeled WT and 3xTg neurons (**Figure 7I**). The ISI histogram was shifted leftward, in line with an elevated firing rate, but did not display a major change in shape (**Figure 7J**).

These simulations suggest that the alterations in intrinsic properties and synaptic inputs measured in 3xTg dopamine neurons are sufficient to produce substantial increases in their firing rates *in vivo* but were not enough to drive the cell into burst firing under basal input. Alternatively, these changes might be homeostatic, acting to maintain firing rates despite an altered balance of presynaptic firing activity in excitatory and inhibitory afferents.

### Predicted impact on burst firing

Because dopamine neurons generate bursts and pauses *in vivo* in response to behaviorally relevant inputs, we performed additional simulations to evoke bursts and pauses. The firing rates of the excitatory inputs were varied from 0 to 20 Hz in 2 Hz increments, while maintaining the inhibitory inputs at 4 Hz. The change in excitation was delivered for 500 ms, to mimic a dopamine neuron burst. When the excitation was halted (0 Hz), a complete cessation of firing was observed in both WT (**Figure 8A,B**) and 3xTg models (**Figure 8C,D**), indicating that baseline synaptic inhibition was sufficient to suppress autonomous pacemaker firing. At the highest excitatory input tested (20 Hz), WT neurons increased firing (**Figure 8E,F**), but the 3xTg group did so to an even greater extent (**Figure 8G,H**).

**Figure 8.**
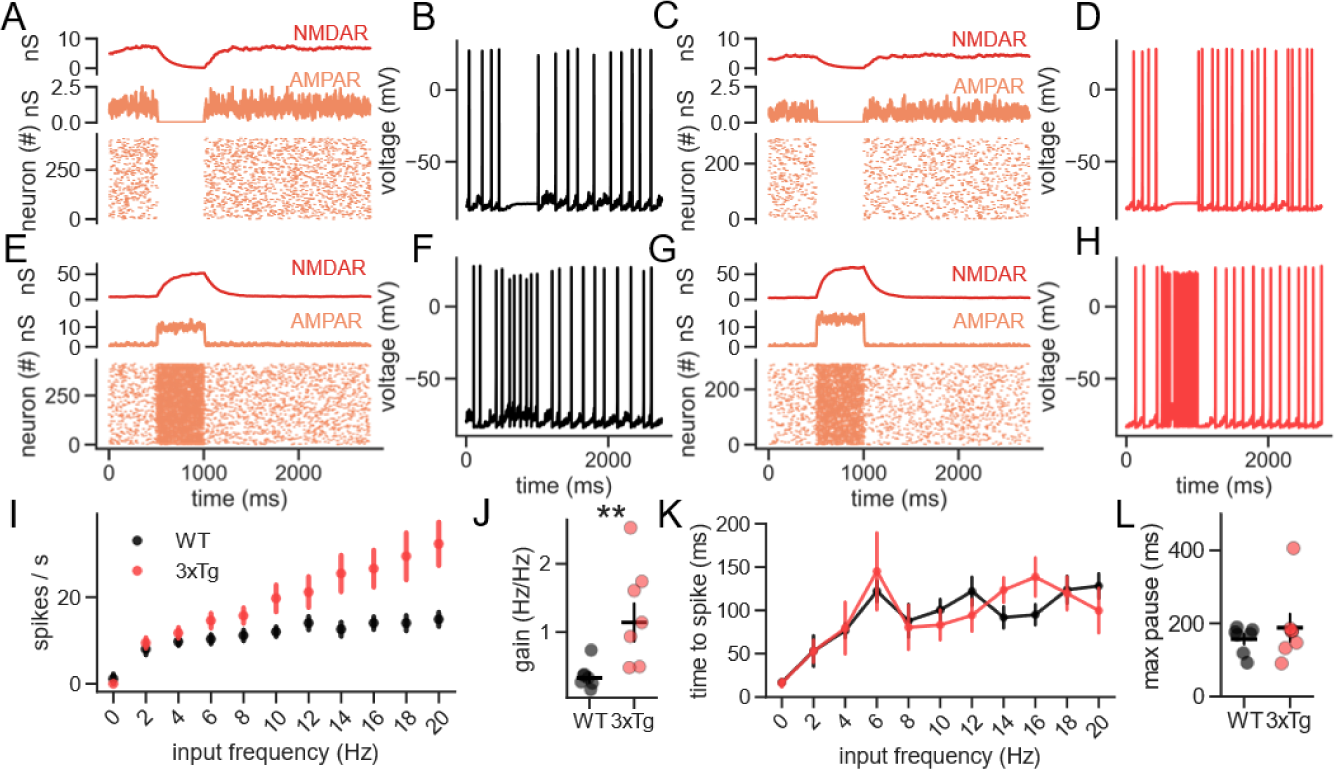
Biophysical modeling predicts that 3xTg neurons are more sensitive to changes in excitatory afferent firing rates. **A**. Simulation of WT dopamine neuron response to a pause in presynaptic excitatory neuron firing. Top trace is the total NMDAR conductance (red). Middle trace is the total AMPAR conductance (orange). Raster plot shows the spike output of the postsynaptic neuron on multiple trials. **B**. Somatic membrane potential response to the pause in synaptic excitation. **C-D**. Same as A-B, but in the 3xTg model. **E-F**. Same as A-B, in a WT model, but for an elevation of presynaptic firing to 20 Hz. **G-H**. Same as E-F, but for a 3xTg model. **I**. Relationship between mean excitatory afferent firing frequency and mean postsynaptic firing rate for WT and 3xTg models (n=8/group). **J**. 3xTg model neurons fire faster at each excitatory input frequency and have a steeper relationship between excitatory afferent frequency and postsynaptic firing frequency (measured from 2-20Hz; Mann-Whitney, U=3.0, P=0.0041). **K.** Relationship between excitatory afferent firing frequency and the pause duration following a burst. **L.** WT and 3xTg models pause similarly following bursts.

To test if increased presynaptic firing rate has differential effects on WT and 3xTg dopamine neuron output firing rates, we plotted output firing frequency against presynaptic input frequency. Across the entire frequency range, the 3xTg group fired more action potentials per second (**Figure 8I**) and had a steeper frequency-frequency relationship (**Figure 8J**). The stronger bursting of model 3xTg neurons was not associated with a longer pause after each burst, as WT and 3xTg models paused for similar durations across the range of presynaptic frequencies (**Figure 8K**) and had similar maximum pause durations (**Figure 8L**).

These results predict that for given firing rates in excitatory and inhibitory afferents, dopamine neurons in 3xTg mice generate action potentials at an elevated rate compared to those in WT mice. In response to barrages of increased excitation, which likely occur during presentation of salient stimuli, VTA dopamine neurons in 3xTg mice may respond more sensitively, generating bursts that release more dopamine from their remaining terminal boutons. Thus, intrinsic and synaptic changes in dopamine neurons may compensate for axonal degeneration in early stages of AD pathology.

## Discussion

Findings from animal models of AD suggest that the impact of amyloid and tau pathology on neuronal activity and synaptic transmission varies widely across brain regions and cell types. Synaptic hypoactivity is commonly reported, likely due to neuronal loss(*92*), and several types of neurons show intrinsic hyperexcitability(*10, 40, 93, 94*). In contrast, reports of elevated synaptic activity are rare(*51, 62, 65*). Here, we show that a model of AD pathology induces morphological and synaptic changes in dopamine neurons, including enhanced synaptic excitation. Biophysical modeling predicts that, together with previously described intrinsic changes, the net result is an elevated basal firing rate and substantially increased input-output gain in response to excitatory afferent firing *in vivo*. These findings provide a plausible mechanism for maintenance of dopaminergic signaling in the face of axonal degeneration(*10, 11*).

Identifying specific pre- and postsynaptic alterations in a chronic disease model can be challenging as, unlike with acute manipulations, the set of synaptic connections itself may vary across groups. Here we showed that the 3xTg model exhibits morphological alterations of dopamine neuron dendrites, which necessarily involves loss or rearrangement of some synaptic inputs and possibly presynaptic axonal arbors as well. Despite the substantial loss of dendritic surface area, 3xTg mice exhibited a clear presynaptic enhancement of glutamate release, evidenced by increased mEPSC frequency, a decrease in PPR, reduced block of evoked EPSCs by kynurenate (suggesting higher synaptic glutamate concentrations), and a smaller relative increase of mEPSC frequency by PDBu. Increased vesicle release is likely a major contributor to higher evoked EPSC amplitudes and an enlarged readily releasable pool, with the caveat that both of these measures may also reflect postsynaptic changes. We also provide evidence of postsynaptic enhancement of AMPA receptor currents, including larger mEPSC amplitudes and higher AMPAR:NMDAR EPSC ratios, although the mechanisms involved are less clear. The absence of LTD after low-frequency stimulation also presumably reflects a postsynaptic change, although the direction of this effect did not correspond to previously reported forms of postsynaptic plasticity in dopamine neurons.

Our measurements of unitary EPSCs indicated that the enhancement of excitatory transmission is not uniform across all synapses but is restricted to a subset of inputs accounting for the top quartile of uEPSC amplitudes. These inputs showed fast rise times, suggesting they form synapses near the soma. Using immunolabeling of filled cells, we did not detect an increased density of GluA1 puncta at perisomatic sites on 3xTg neurons, arguing against the notion that the strengthened afferents form more synapses. However, we detected a strong negative correlation in the AMPAR puncta intensity as function of distance from somatic center, which may indicate that distal connectivity is inherently weak in WT and 3xTg mice, alike. Therefore, loss of distal dendrites may have relatively little impact on total synaptic excitation. AMPA currents in 3xTg mice did not exhibit altered rectification, which would have also been a clear indication of postsynaptic alterations. Rather, the affected synapses are likely the main sites of presynaptic potentiation, with the apparent postsynaptic changes reflecting the enhanced weight of connections that feature different baseline properties (e.g., mEPSC amplitudes, AMPAR:NMDAR ratio, and susceptibility to LTD). In future studies, it will be important to identify the source of these strengthened inputs using optogenetic methods to characterize the synaptic changes with greater specificity and to understand their impact on circuit function.

While the molecular mechanisms of excitatory synaptic enhancement remain under investigation, the partial occlusion of PDBu enhancement of mEPSCs and the recovery of mEPSC frequency with PKC inhibition could implicate activation of presynaptic PKC as a driving mechanism in 3xTg mice. In other neuron types, presynaptic PKC activation has been associated with enlargement of the RRP and a decreased energy barrier for release, leading to increased release probability(*59, 66, 67*), with downstream targets that include liprin-α3, synaptotagmin-1, synaptobrevin, and SNAP-25(*61, 73, 95–97*). Our results suggest that baseline phosphorylation of PKC targets might be greater in 3xTg neurons. Several studies have found that elevated PKC activity alters synaptic transmission in AD models(*62–64*), and PKC has been implicated in pre- and postsynaptic mechanisms of LTP in VTA dopamine neurons(*70*). For these reasons, PKC-related signaling cascades are of major interest for future study and potential intervention. However, because PKC is a ubiquitous kinase with numerous targets in the central nervous system(*98*) and elsewhere, establishing specificity will be essential.

Previous studies show that dopamine neurons can exhibit long-term postsynaptic enhancement of excitatory input by increasing surface expression of GluA2-lacking, calcium-permeable AMPA receptors(*77, 99, 100*). This canonical form of potentiation also enhances LTD induced by a sustained 1 Hz field stimulation, indicating that the potentiation is reversible. In contrast, 3xTg dopamine neurons did not show inward-rectifying AMPAR currents, inconsistent with the addition of GluA2-lacking AMPA receptors. We also observed a loss of LTD after low-frequency stimulation in 3xTg neurons, suggesting a possible postsynaptic change in AMPAR membrane localization. Postsynaptic enhancement and diminished plasticity may reflect an imbalance in AMPAR turnover, as a recent study that investigated the role of the AAA+-ATPase Thorase, which modulates AMPAR internalization, reported similar findings(*79*). In a previous Patch-seq screen of dopamine neurons in WT and 3xTg mice, we did not detect changes in the Thorase-encoding gene *ATAD1*(*10*); however, microtubule destabilization might alter surface receptor trafficking, producing the observed effects. It is possible that the loss of excitatory synaptic plasticity may reflect an inability to adapt to changing environmental cues, and may contribute to reduced reward learning behavior that we previously reported in 12-month-old 3xTg mice(*10*).

The synaptic alterations in VTA dopamine neurons from 3xTg mice were accompanied by a profound reduction in somatodendritic arbor and surface area. One possible model-driven mechanism for this is the hyperphosphorylation of tau, resulting in destabilization of microtubules. In AD patients, synaptic loss is tied to both tau phosphorylation and network connectivity(*101*). Similar changes in morphology were reported previously in SNc dopamine neurons in the metabolic MitoPark and MCI-Park models of Parkinson’s disease (*41, 42*) but not in healthy aged mice(*43*). Somatodendritic retraction might therefore be a conserved homeostatic mechanism in diverse diseases affecting dopamine neuron function and viability. Not only would dendritic retraction decrease the cell volume to be maintained in the face of the building pathology, but decreased surface area could enhance the efficacy of remaining synaptic inputs by reducing the electrotonic distance to the axonal spike initiation zone. In dopamine neurons recorded *in vivo*, there is no correlation between total dendritic length and firing rate or variability(*102*), suggesting that homeostatic adaptations of synaptic input and/or intrinsic ionic mechanisms can normally maintain consistent spike output among neurons of varied morphology. However, these recordings were conducted in anesthetized mice, and different results might be found in awake animals, where excitatory and inhibitory input are likely significantly greater(*103*). We also observed a modest reduction in GABA_A_R-mediated IPSCs in 3xTg mice, but no reduction in unitary IPSC amplitudes, which could reasonably be explained by a frank loss of synapses on a pruned dendritic arbor. Interestingly, SNc dopamine neurons in the MCI-Park model of Parkinson’s show *enhanced* GABA_A_ transmission(*42*), arguing against a uniform set of synaptic consequences to dendritic retraction in dopamine neurons.

Combining our measurements of synaptic responses, intrinsic firing properties, and cell morphology in compartmental biophysical models of VTA dopamine neurons, we predict that neurons in the 3xTg mice respond more sensitively to changes in presynaptic activity, particularly for excitatory afferents. At present, it is not known whether this increase in response gain is pathological or homeostatic. The change may compensate for decreased projection region innervation reported previously(*10, 11, 16*), helping to maintain phasic dopamine signaling required for reward learning. However, this compensation may not be enough to completely maintain reward learning in 3xTg mice(*10*). Alternatively, the increased gain of 3xTg neurons might alter the pattern of dopaminergic output in a manner that is detrimental for their projection targets. *In vivo* manipulations of WT and 3xTg dopamine neuron activity would be required to determine the behavioral consequences of altered neuronal function in this model.

## Materials and Methods

### Mice

All animal experiments were performed in compliance with Oklahoma Medical Research Foundation’s Institutional Animal Care and Use policies. WT and 3xTg-AD founders (MMRRC #034830, on the C57BL/6;129×1/SvJ;129S1/Sv embryo injected in a B6;129 hybrid background) were originally acquired from Dr. Salvatore Oddo(*39*) and were bred to homozygosity for all three transgenes (APP_Swe_, PSEN1_M146V_, and MAPT_P301L_) in-house. Male and female mice were housed on a reverse light cycle (12 hours light, 12 hours dark, lights off at 0900). *Ad libitum* access to food and water was provided to all mice. Breeder genotyping was outsourced to Transnetyx. Both sexes were used in each experiment in roughly equal numbers, and the data for male and female mice were pooled.

### Brain slice preparation

Sections for morphological assessment, immunohistochemistry, and patch clamp electrophysiology were collected as described previously(*10*). First, mice were deeply anesthetized with isoflurane and decapitated. The brain was rapidly removed and placed in ice-cold cutting solution containing the following (in mM): 110 choline chloride, 2.5 KCl, 1.25 NaH_2_PO_4_, 0.5 CaCl_2_, 10 MgSO_4_, 25 glucose, 11.6 Na-ascorbate, 3.1 Na-pyruvate, 26 NaHCO_3_, 12 N-acetyl-L-cysteine, and 2 kynurenic acid. 200 µm horizontal sections containing the ventral midbrain at the level of the *fasiculus retroflexus* (fr) and medial terminal nucleus of the accessory optic tract (*mt*) were cut on a VT1200S vibrating microtome (Leica). Slices were then transferred to an incubation chamber at 32 °C containing (in mM) 126 NaCl, 2.5 KCl, 1.2 MgCl_2_, 2.4 CaCl_2_, 1.2 NaH_2_PO_4_, 21.4 NaHCO_3_, and 11.1 glucose for thirty minutes before transferring to room temperature for at least thirty additional minutes prior to the beginning of the experiment. For experiments not involving NMDA currents, MK-801 was added to the recovery chamber to block NMDARs, to minimize excitotoxicity.

### Neuronal reconstruction and analysis

Single dopamine neurons were dialyzed with 5 mg/mL biocytin through a recording pipette for at least five minutes. Slices containing filled neurons were fixed overnight in 4% paraformaldehyde, permeabilized, and stained against using streptavidin conjugated to a fluorophore (1:500, Invitrogen streptavidin-647, #S21374). Brain slices were cleared using AKS(*104*) and imaged on a Zeiss LSM 880 through the entirety of the horizontal section (200 µm). Individual neurons were traced using the Simple Neurite Tracer(*105*) plugin in FIJI(*106*) in 3D to generate .TRACE file skeletons. Three-dimensional skeletons were used to compute the Sholl analysis with the built-in Simple Neurite Tracer function. After tracing, a binary mask was generated using the fill function to generate a .tiff file containing the neuron Boolean mask. This mask was then imported in Python using aicsimageio, and the marching cubes scikit-image function was used to extract the 3D surface mesh from the 3D volume in the biocytin channel. This functions by computing a polygonal mesh from the surface of the neuron, creating a model of each neuron, with high-fidelity surface detection due to the small cube size. The model can then be used to calculate the surface area of each cell using skimage.measure.mesh_surface_area, from which the polygonal face values can be multiplied by voxel size (420 nm) to compute the cellular surface area. Cellular volume was computed by counting all voxels within the surface mesh and multiplying by voxel volume. All skeletonized reconstructions can be seen in **Figure S1** and will be submitted to NeuroMorpho.org upon acceptance of the manuscript. Three-dimensional morphological depictions of dendrites in **Figure S7G,H** were created in Blender from surface meshes of the biocytin mask and AMPAR puncta.

### Patch clamp electrophysiology

Acute brain slices were transferred to a recording chamber and perfused with warmed aCSF (31-33°C) using an inline heater (Warner Instruments) and a peristaltic pump (Warner Instruments) at a rate of ∼2 mL/min. Dödt gradient contrast was employed to visualize the tissue on an upright microscope (Nikon). Dopamine neurons of the VTA were identified anatomically and by electrophysiological parameters, as described previously(*10*). We selected dopamine neurons largely in the parabrachial pigmented nucleus (PBN) containing cell bodies located near the *fr* and >100 µm from *mt.* Dopamine neuron identity was confirmed by the well-characterized physiological parameters of this cell type. Notably, VTA dopamine neurons display slow (0.5-5 Hz), rhythmic, spontaneous firing(*90*) governed by a slow A-type potassium conductance (inactivation time constants between 30 and 300 ms)(*86, 91*). These properties are distinct from those of local GABAergic neurons(*107–109*). Patch pipettes were pulled from thin-walled glass (Warner Instruments) and had resistances of 2.5-3.5 MΩ. Intracellular solutions for GABA_A_ receptor mediated responses contained (in mM) 140.5 CsCl, 5 Qx314-Br, 7.5 NaCl, 10 HEPES, 0.1 EGTA, 2 Mg-ATP, and 0.21 Na-GTP, pH adjusted to 7.35 with CsOH. For AMPA and NMDA receptor mediated responses, the same internal solution was used except that CsCl was replaced with equimolar Cs-methylsulfonate. The liquid junction potential (-11 mV for Cs-methylsulfonate and -5 mV for CsCl) was not corrected unless otherwise noted. Series resistance was measured from the current transient produced by a small hyperpolarizing voltage step, and recordings were discarded if series resistance exceeded 15 MΩ or varied by more than 25% during a recording. For morphological reconstructions, Cs-methylsulfonate internal solution was used with the addition of 5 mg/mL biocytin. All recordings were obtained using AxoGraph version 1.8.0, and data were low-pass filtered at 6 kHz and sampled at 20 kHz. The holding potential for all recordings was -60 mV with the exception of AMPAR:NMDAR ratio and NMDA bath application experiments, which were conducted at +40 mV to relieve voltage-dependent magnesium block.

### Electrical stimulation

Biphasic current pulses (0.1 ms) were produced by an A365 stimulus isolator (World Precision Instruments) and delivered via a platinum-coated bipolar stimulating electrode (FHC) placed in the caudal VTA. The stimulation intensity used for eEPSCs in the RRP train analysis was 2x threshold. The stimuli used for IPSC PPR and amplitude determination were 100 µA. Minimal stimulation of unitary evoked currents was conducted as described previously(*84*), by manual adjustment of stimulus intensity to generate visible failures (no PSC) and successes at roughly a 1:1 ratio.

### Western blot

Mice were euthanized by decapitation under isoflurane-induced anesthesia and brains removed rapidly and submerged in ice cold cutting solution. Brains were then placed in a 1mm brain block (Alto) and a section containing the ventral midbrain was extracted (approximately Bregma -3.6 to -2.6, **Figure S3A**). From the extracted slice, the ventral midbrain was further dissected (**Figure S3B**). Each sample was comprised of two of these ventral midbrain subsections, one from each of two mice, in 200 µL RIPA buffer (Nalgene) with 2% Halt protease and phosphatase inhibitor (Thermo Scientific). Sections were homogenized, then centrifuged at 2000×g and 4°C for 8 minutes. The supernatant was used as a working lysate, to which lithium dodecyl sulfate sample buffer and dithiothreitol were added prior to heating at 95°C for 5 minutes. For western blot analysis, samples were run on 4-12% Bis-Tris gradient gels (Invitrogen) at 200V for 45 minutes and transferred to nitrocellulose membranes at 30V for 1 hour using a semi-dry transfer system (Novex, Life Technologies). Total protein was stained using Revert 700 Total Protein Stain (LI-COR) and imaged using Odyssey CLx (LI-COR) for later normalization. Blots were incubated overnight at 4°C in primary antibodies (**Table S2**) at 1:500 in blocking buffer (LI-COR) with 1% tween, then washed 3 times for 10 minutes each in tris-buffered saline with 1% tween (TBS-T) and incubated for 1 hour at room temperature in secondary antibody at 1:2000 (Goat anti-Rabbit, 926-32211, LI-COR). Following 2 washes (10 minutes each) in TBS-T and one wash in TBS, blots were again imaged using the LI-COR system and quantified using Image Studio (Version 5.2).

### Electrophysiology analysis

Miniature synaptic currents were detected by first smoothing currents via convolution with a Gaussian waveform (0.4 ms standard deviation). The derivative of the smoothed trace was taken, and negative peaks in the derivative were detected with a threshold of -3.5 times the interquartile difference. The false positive rate for mEPSC detection was < 0.1 event per second, assayed by DNQX wash-on, and was robust to changes in mEPSC amplitude (**Figure S2**). The rectification index was corrected for liquid junction potential and was calculated as 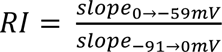. Unitary synaptic currents were smoothed lightly by Gaussian convolution (0.1 ms SD), and successes and failures were identified by manual inspection of each waveform. In experiments producing more than one distinct unitary response identifiable by latency, amplitude, and/or waveform, each group of responses was identified and analyzed separately. Recordings showing signs of multiple unitary responses that were not readily separated were rejected. All electrophysiology data analysis was conducted in Python, and code is available on the corresponding author’s GitHub (https://github.com/heblanke/Elevated_excitation_dopamine_neurons_3xTg/).

### Immunohistochemistry

Following biocytin filling during patch clamp recordings, slices were transferred to 4% PFA overnight, washed in PBS, and stored at -20°C in cryoprotectant until use. All brain sections were processed in parallel to minimize experimental artifacts. Sections were washed in PBS to remove any excess cryoprotectant. 5 mg/mL NaBH_4_ in PBS was used to quench peroxidase activity and minimize background. Slices were permeabilized following the iDISCO protocol(*110*) in a solution containing 80% (v/v) PBST, 20% (v/v) DMSO, and 2.3% (w/v) glycine, in order to improve antibody penetration. Sections were blocked in 5% bovine serum albumin and 0.5% normal donkey serum overnight. Slices were again washed in PBST and incubated with primary antibodies (guinea pig anti-GluR1 Alomone [AGC-004-GP], 1:250) and were then incubated with streptavidin-647 (1:500, Jackson Immuno, 016-600-084) and anti-guinea pig Alexa Fluor 568 (1:500, Invitrogen A-11075) for 48 hours. Sections were washed, then cleared using the AKS protocol (20% (v/v) TDE, 20% (v/v) DMSO, 20% (w/v) sorbitol, and 0.5 M Tris base, in H_2_O)(*104*). The AKS tissue clearing minimized light scatter and prevented substantial intensity loss as a function of sample depth. The sections were then mounted on charged glass slides with a 200 µm agar spacer, dried slightly, and coverslipped with AKS clearing solution.

### Confocal microscopy

Images were acquired on a Zeiss 880 laser scanning microscope with Zen Black software. Individual dopamine neuron somata were easily identified by streptavidin fluorescence. To capture the entire somatodendritic tree, Z-stack tiled images including the entire neuron were acquired using a water-immersion 40x objective. Identical settings were used for every image acquisition to allow for comparison of labeling intensity between groups.

### Detection of AMPA receptor puncta

Binary masks generated using the fill method in SimpleNeuriteTracer were imported into Python using AICSImage (https://github.com/AllenCellModeling/aicsimageio), along with the GluA1 channel of the image data. For detection, 80% of the maximum intensity values for each Z-slice were computed, and a normalized Z-stack was generated from the maximum and minimum values. Using the normalized Z-stack, we applied a Richardson-Lucy deconvolution, which uses a probabilistic approach based on Poisson statistics, using the point spread function over ten iterations. This deconvolution method optimized the relationship between sharpness and development of artifacts. To remove background and detect AMPAR puncta, we determined the interquartile range of values in the GluA1 channel within the biocytin mask, and computed a threshold value as follows: *Threshold* = *Q*75 + 1.5 ∗ (*Q*75 − *Q*25), where Q75 is the 75^th^ percentile and Q25 is the 25^th^ percentile, and set all values below this threshold to zero, which strongly resolved the GluA1 puncta. The image arrays were then converted to Booleans using Sauvola local thresholding implemented in SciPy filters. GluA1 puncta were then localized and counted using skiimage.measure.regionprops, which determines the coordinates and size of the puncta.

### Distance from the soma

To determine the center of the soma algorithmically, we used the biocytin binary mask and used a distance transformation implemented in ndimage (distance_transform_edt) to find Euclidean distances that maximized the minimum distance to the edge of the biocytin mask. From the 3D mask, we skeletonized each neuron using skiimage.morphology and converted that skeleton into a networkX graph in Python. The graphs were used to calculate intensity values as a function of distance from cell soma via networkX built in function single_source_dijkstra_path_length, which computes the shortest distance between points along each branch. The maximum distance from the soma varied among cells. For subsequent analysis, these values were rescaled between 0 and 1 (**Figure S7I**), and the slope between rescaled distance and GluA1 intensity was calculated (**Figure S7J**).

### Quantification of GluA1 puncta

Following GluA1 detection, the puncta masks were applied to the original image. Values of each individual mask were collected and stored using skiimage.measure.regionprops.

### Biophysical modeling

Multi-compartment biophysical models were created using Jaxley(*111*) in Python. TRACE files generated from SimpleNeuriteTracer were converted to SWC files using *xyz2swc’*s web-based converter (https://neuromorpho.org/xyz2swc/ui/)(112). SWC files were then imported into Jaxley’s custom SWC reader using the graph-backend, which relies on networkX graphs, to allow for morphological manipulation and attachment of a spike-generating axon segment(*111*). Each dendritic branch was comprised of four compartments. The ionic conductances included a voltage-gated eight-state Markov Na+ conductance, a non-inactivating Na+ conductance, a five-state voltage-gated delayed-rectifying K+ conductance, an A-type potassium conductance, a calcium-activated SK-type potassium conductance, a leak K+ conductance, an M-type K+ conductance, an L-type Ca+ conductance, an N-type Ca+ conductance, and a Ca+ pump. The ionic conductance values were allowed to vary around values previously implemented in biophysical models of dopamine neurons(*89, 113, 114*), and were tuned using stochastic gradient descent(*111*) on an archetypal WT neuron, arriving at a WT cell state (**Table S1**). These conductance values were applied to every neuronal morphology to minimize additional confounds. A multi-compartment axon segment was added, branching from the soma, to act as the spike generator. The axon had sodium channel conductance values higher than those of the other compartments to ensure that this segment was the site of action potential initiation. In the 3xTg neuron models, the SK conductance was lowered by 50% to increase intrinsic excitability in line with our previous report(*10*).

Excitatory and inhibitory afferents were randomly connected to the model neuron, with a connection probability proportional to the surface area of each compartment. The amplitudes and time constants of the excitatory and inhibitory conductances were based on our unitary EPSC and IPSC data but did not include quantal variance or synaptic failures. The firing activity of each afferent was simulated as a Poisson process using the Poisson generator in numpy. At baseline, each excitatory afferent was activated at an average rate of 2 Hz. Thus, a WT dopamine neuron model of average surface area received excitatory inputs at a total rate of 1000 Hz. Inhibitory afferents were activated at an average rate of 4 Hz, providing inhibitory inputs at a total rate of 2000 Hz to the average WT neuron. Each presynaptic spike generated a postsynaptic conductance in one compartment of the model neuron (**Figure S11**). In the 3xTg neuron models, the perisomatic excitatory inputs (those arriving on the soma or on a first-order dendrite) were scaled to four times the conductance amplitude of the other excitatory inputs. The conductance parameters of the models are listed in **Table S1**. For each simulation, an initialization period of at least 250 milliseconds was not analyzed to allow for equilibration of model states. All code related to the model is deposited on the corresponding author’s GitHub (https://github.com/heblanke/Elevated_excitation_dopamine_neurons_3xTg/), and upon acceptance of the manuscript, all model details will be submitted to model.db.

### Statistical analysis

Sample size was not predetermined by any statistical methods. Normality of data was assessed by a Shapiro-Wilk normality test (*P* < 0.05 indicating a non-normal distribution). A two-tailed Student’s *t* test was used for pairwise comparisons, and two-way analysis of variance (ANOVA) followed by Sidak’s post hoc test was used for four groups, as indicated in the figure legends. For data that were not normally distributed, nonparametric alternatives, such as Mann-Whitney or Kruskal-Wallis tests, were used. All data in bar graphs and summary plots were shown as means ± SEM, unless otherwise noted in the figure legend. Cumulative probabilities were assessed with either Kolmogorov-Smirnov or Singleton-Epps tests. Significance was indicated by \**P* < 0.05, \*\**P* < 0.01, \*\*\**P* < 0.001, and \*\*\*\**P* < 0.0001. All statistical analyses were performed in Python 3.9 using the scipy.stats package.

## Supporting information

All supplemental materials

## Acknowledgments

We thank Kelsey Carter and Nicole Yates for their excellent technical support. We would like to thank the OMRF Imaging Core, especially Ben Fowler and Julie Crane, for their assistance imaging single, reconstructed neurons. We are immensely grateful to Alexander Lin and Kylie Handa, who worked to troubleshoot dopamine neuron filling, imaging, and reconstruction. We would like to also thank Dr. Holly Van Remmen for sharing research materials.

## Author contributions

Conceptualization: HEB

Methodology: HEB, MHH, KyMH

Investigation: HEB, MHH, KyMH

Analysis: HEB, MHH, KyMH

Visualization: HEB, MHH, KyMH

Writing (original draft): HEB

Writing (review and editing): HEB, MHH, KyMH, KMH, MJB

Funding acquisition: HEB, KyMH, KMH, MJB

Supervision: MJB, KMH

## Data and materials availability

All data are available in the main text or the supplementary materials. Raw data files are available upon request.

