## Supplementary material for "PKC-dependent enhancement of glutamate input to VTA dopamine neurons in 3xTg-AD mice": All supplemental materials

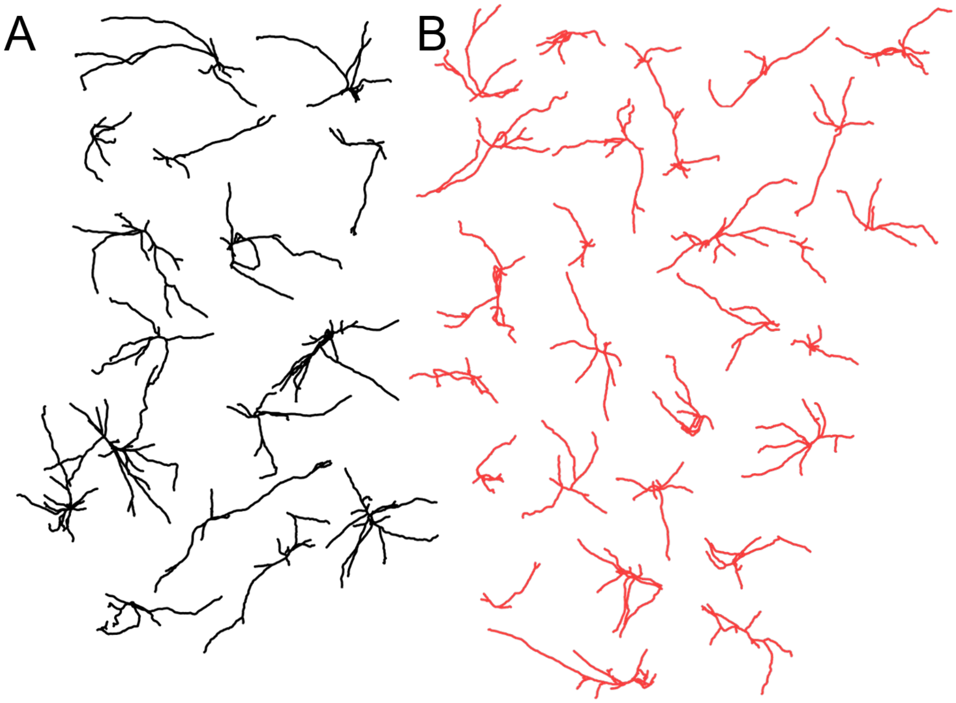


**Supplemental Figure 1. Dopamine neurons in WT and 3xTg mice display a range of neuronal architectures**. Flattened 2-D schematics of all analyzed WT (**A**, black) and 3xTg dopamine neurons (**B**, red).

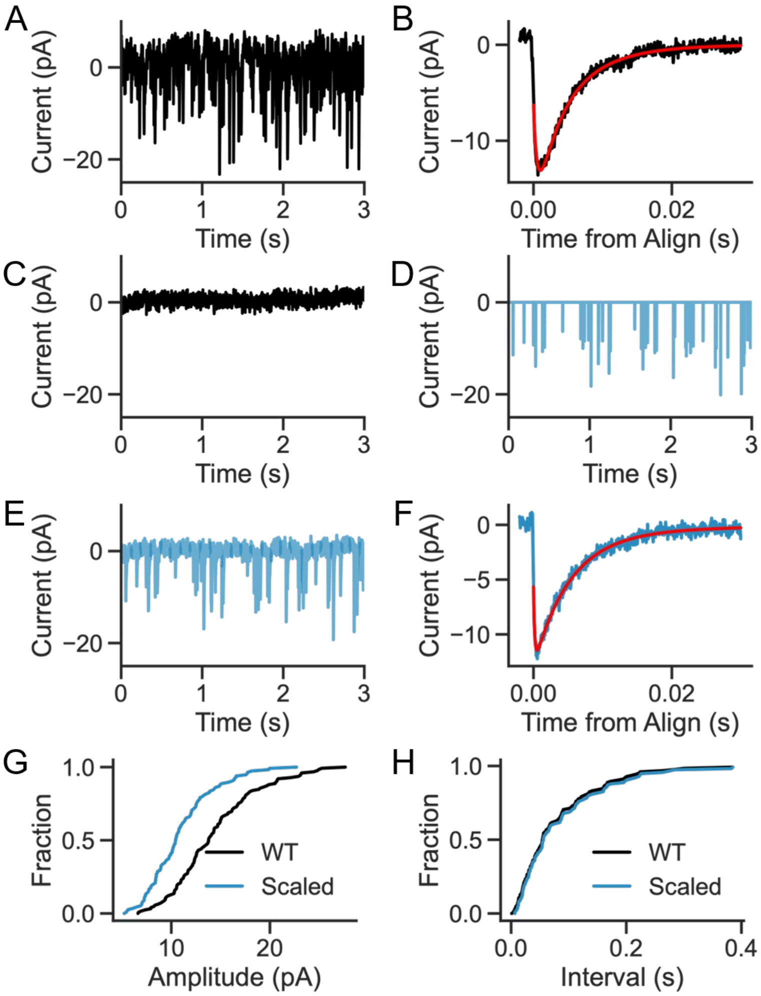


**Supplemental Figure 2. Miniature current detector is robust to changes in amplitude**. To confirm that the increase in mEPSC frequency in 3xTg mice was not an artifact of increased detectability due to larger events, we re-analyzed recorded events using down-scaled amplitudes. **A.** Representative three-second trace of mEPSCs in a PDBu-treated WT dopamine neuron. **B.** Extracted and averaged mEPSC fit with a biexponential curve. **C.** A five second trace following bath application of the AMPA receptor blocker DNQX (10µM). **D.** Event times and amplitudes represented as delta pulses extracted and scaled by the mean difference between WT and 3xTg amplitudes (79.6%). **E.** Point representation convolved with the mean mEPSC bi-exponential waveform and added to the noise from (**C.**). **F.** Modeled, downscaled mEPSCs detected, averaged, and fit. **G.** Cumulative probability of mEPSC amplitudes from WT and downscaled, reconstructed mEPSC data, showing significantly different amplitude distributions (Kolmogorov-Smirnov test, KS statistic=0.3891, P=1.05x10$^{-8}$). **H.** mEPSC down-scaling produced no discernable difference in detected inter-event intervals (KS statistic= 0.0463, P= 0.9985).

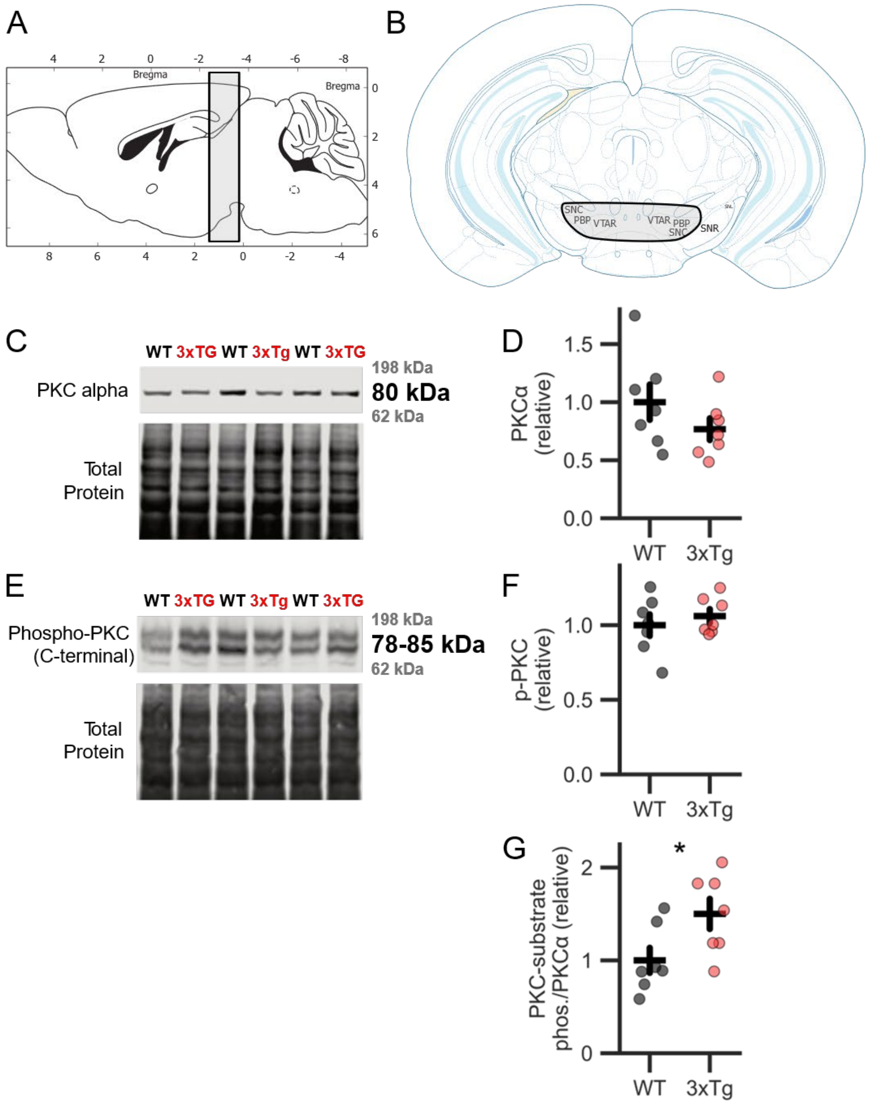


**Supplemental Figure 3. PKCα and phospho-PKC levels in 12 mo WT and 3xTg midbrains. A.** Sagittal representation of 1mm section grossly dissected using a brain-block. **B.** Microdissected ventral midbrain region used for Western blot. **C,D.** Total PKCα levels are not different between genotypes (two-tailed t-test, t = 1.3, P=0.22; n=14 mice, 7 replicates/group). **E**,**F.** C-terminal pan-PKC phosphorylation is also not affected by genotype (two-tailed t-test, t=-0.71, P=0.49). **G.** Phosphorylated PKC substrate is significantly higher relative to PKCα in the 3xTg lysate (two-tailed t-test, t = -2.73, P = 0.035). Figure partially adapted from Paxinos and Franklin, 2019.

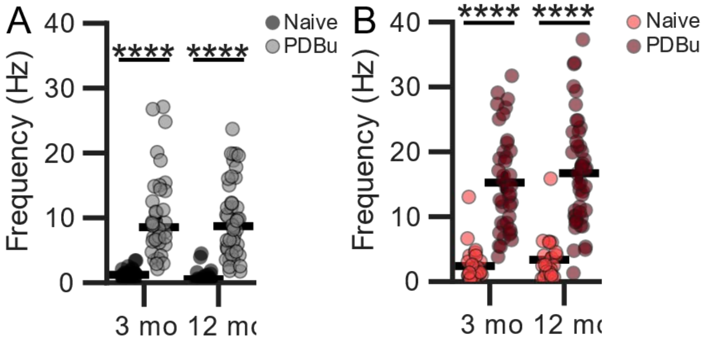


**Supplemental Figure 4. PDBu dramatically enhances mEPSC frequency in WT and 3xTg neurons at 3 and 12 mo. A**.PDBu enhanced the mEPSC frequency in WT neurons (2-way ANOVA main effect of treatment F=123.85, P<1x10$^{-20}$) in both 3 mo (Sidak post-hoc, P=1.09x10$^{-12}$) and 12 mo (Sidak post-hoc, P=5.91x10$^{-10}$) neurons. **B.** PDBu also enhanced mEPSC frequency in 3xTg neurons (2-way ANOVA main effect of treatment F=124.44, P<1x10$^{-20}$) in 3 mo (Sidak post-hoc, P=1.07x10$^{-13}$) and 12 mo (Sidak post-hoc, P=2.65x10$^{-13}$) neurons. Sample sizes, untreated 3 mo (35) and 12 mo (36); treated 3 mo (38) and 12 mo (44).

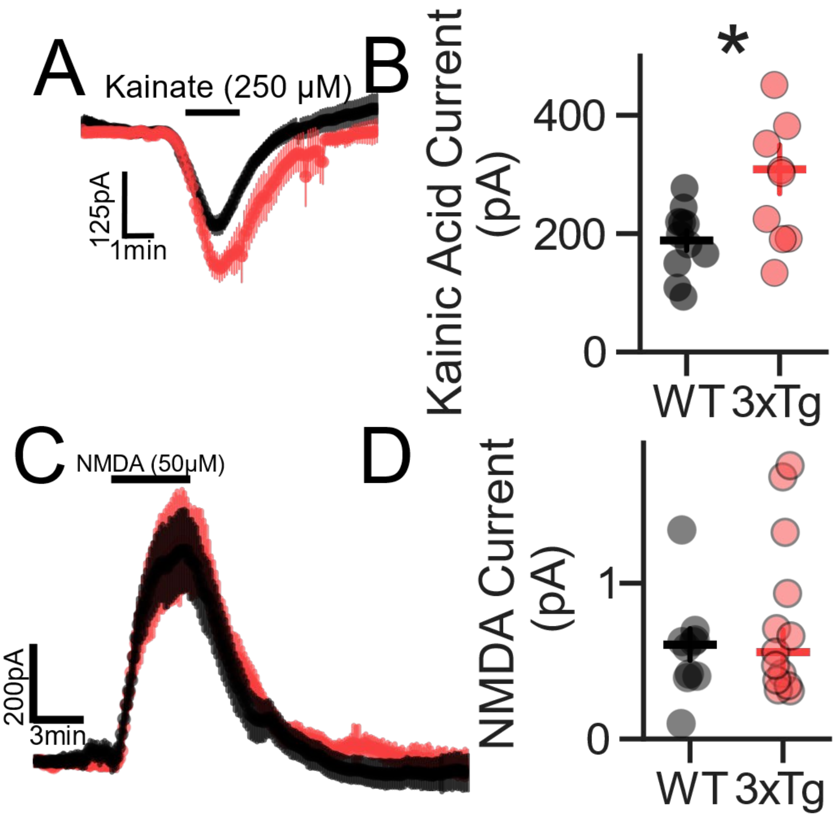


**Supplemental Figure 5. 3xTg dopamine neurons display enhanced sensitivity to kainic acid**. **A**. Inward current produced by bath application of kainic acid (250 µM) in WT (black) and 3xTg (red) neurons. **B**. Maximum current induced by kainic acid is significantly larger in 3xTg dopamine neurons (t=-2.71, P=0.019; n=11 [WT] and 10 [3xTg]). **C**. Outward current produced by bath application of NMDA to WT neurons (black) and 3xTg (red). **D**. WT and 3xTg dopamine neurons displayed a similar maximal response to NMDA (n=10 [WT] and 13 [3xTg]).

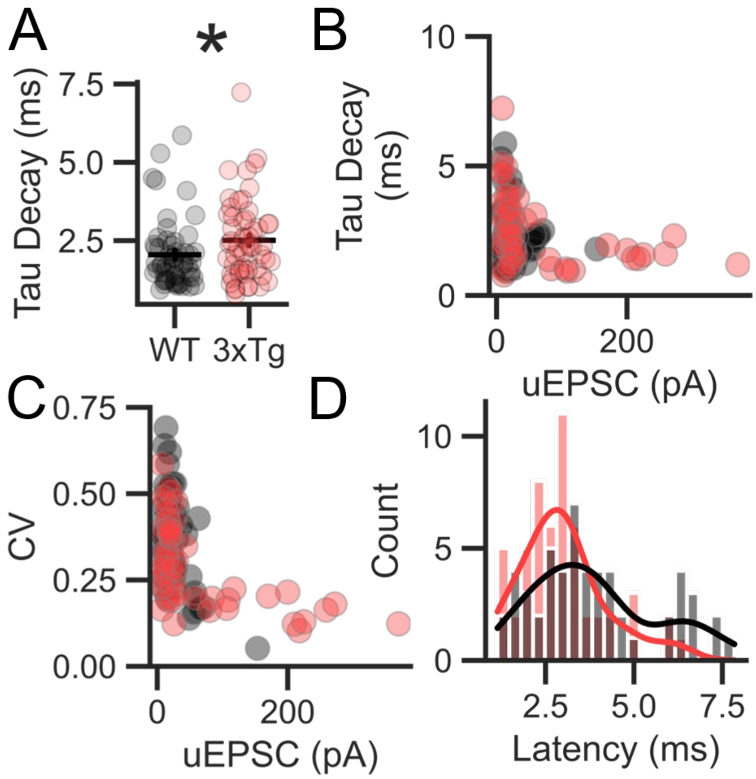


**Supplemental Figure 6. uEPSC kinetics and CV suggest pre- and postsynaptic enhancement. A.** uEPSC tau decay times are slower in the 3xTg than in WT dopamine neurons (Mann-Whitney, P=0.032, U=1839.0; n=57 [WT] and 71 [3xTg]), suggesting a change in AMPA receptor properties. **B.** The largest amplitude uEPSCs had the fastest decay time constants, consistent with a proximal location of these inputs. **C.** The largest uEPSCs had the lowest coefficients of variation of their amplitudes (measured for successes only), suggesting a larger quantal content, compared to smaller uEPSCs. **D.** The latency from the stimulus was similar between WT and 3xTg uEPSCs.

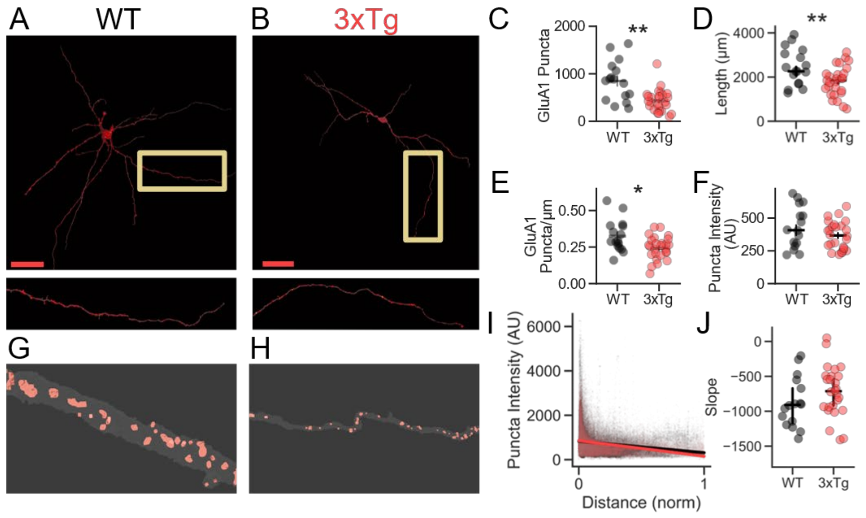


**Supplemental Figure 7. AMPA receptor expression is not increased in 3xTg dopamine neurons**. **A**. Representative WT dopamine neuron with masked volume defined by biocytin-streptavidin labeling (gray) and GluA1 staining (red). Top panel is a maximum-intensity projection in the x-y plane, and bottom panel is an enlarged image of the yellow-boxed segment. **B**. 3xTg dopamine neuron. **C**. The total number of GluA1-positive puncta was lower in 3xTg dopamine neurons compared to WT neurons (Mann-Whitney, P=0.0013, U=333.0; n=13 [WT] and 17 [3xTg]). **D**. The total length of 3xTg dopamine neurons was lower than WT (t=2.82, P=0.007). **E**. 3xTg dopamine neurons have fewer GluA1 puncta per µm (Mann-Whitney, P=0.0165, U=301.0). **F**. The average intensity of individual GluA1 puncta is similar between WT and 3xTg dopamine neurons. **G**. Three-dimensional reconstruction of a WT dopamine neuron segment showing the biocytin mask in gray and the GluA1 puncta in red. **H**. Same as in G for 3xTg dendritic segment. **I**. GluA1 puncta intensity plotted against normalized distance from soma for all puncta in WT and 3xTg dopamine neurons. **J**. Both WT and 3xTg neurons showed a decrease in GluA1 puncta intensity with normalized distance from the soma, suggesting that more distal synapses had fewer AMPA receptors. The relationship was similar in WT and 3xTg neurons, as calculated by the fitted slope of GluA1 intensity versus distance from the soma.

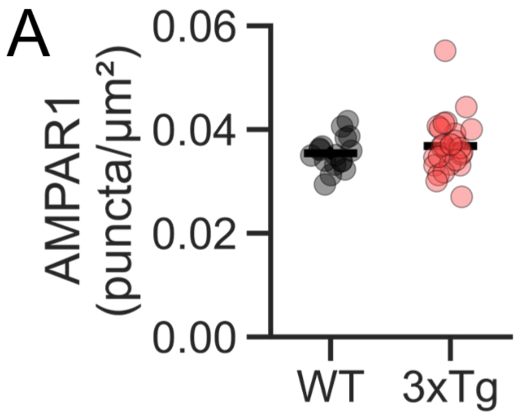


**Supplemental Figure 8. GluA1 puncta densities normalized by surface area. A.** GluA1 puncta densities (puncta/µm2) are similar in WT and 3xTg dopamine neurons (n=13 [WT] and 17 [3xTg]).

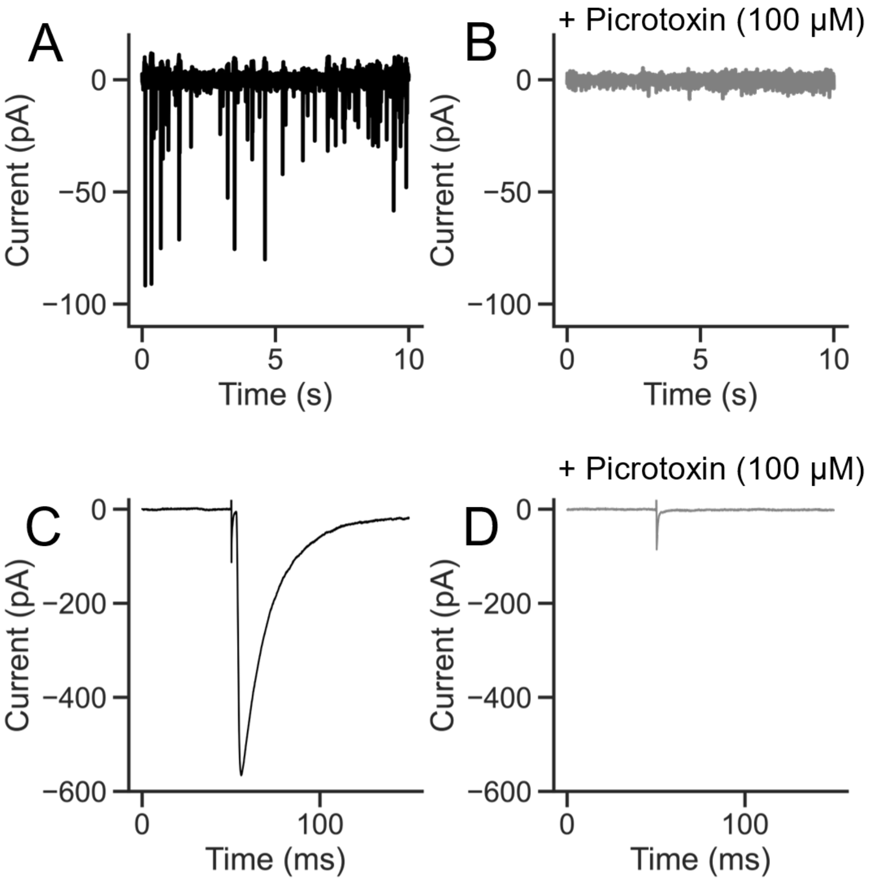


**Supplemental Figure 9. mIPSCs and eIPSCs are mediated by GABA$_A$ receptors. A.** mIPSCs recorded from a VTA dopamine neuron. **B.** Absence of detectable mIPSCs during application of 100 µM picrotoxin. **C.** Average evoked IPSCs (eIPSCs) blocked by picrotoxin (**D**).

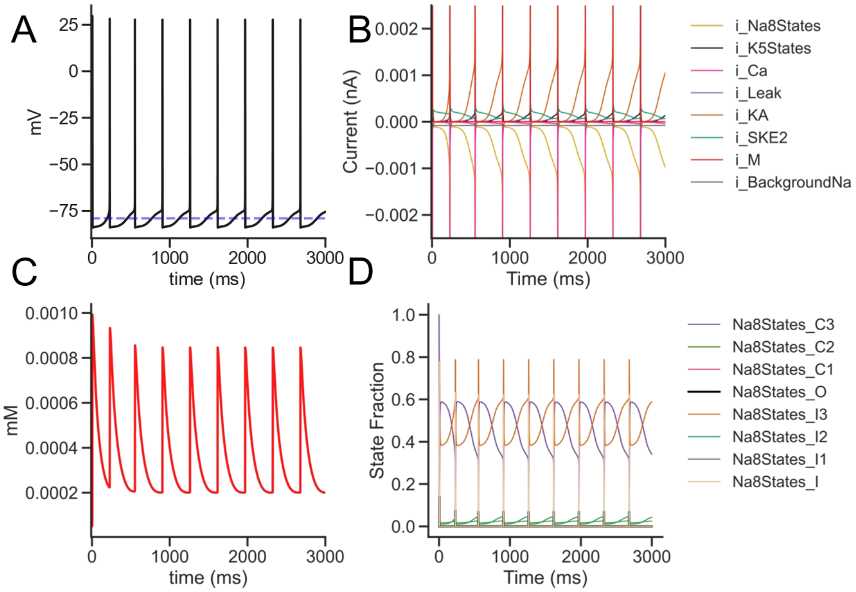


**Supplemental Figure 10. Compartmental biophysical model recapitulates firing rates and currents generated by dopamine neurons in *ex vivo* preparations. A.** Firing of model WT dopamine neuron with detected spike peaks marked in red and the chloride reversal potential indicated by the blue dashed line. **B**. Intrinsic currents during pacemaker firing. Maximum currents for fast sodium and delayed rectifier potassium channels are truncated. **C**. Calcium concentration fluctuation (in mM) during pacemaker firing as measured at the soma. **D**. The Markov-model sodium channel states during pacemaking.

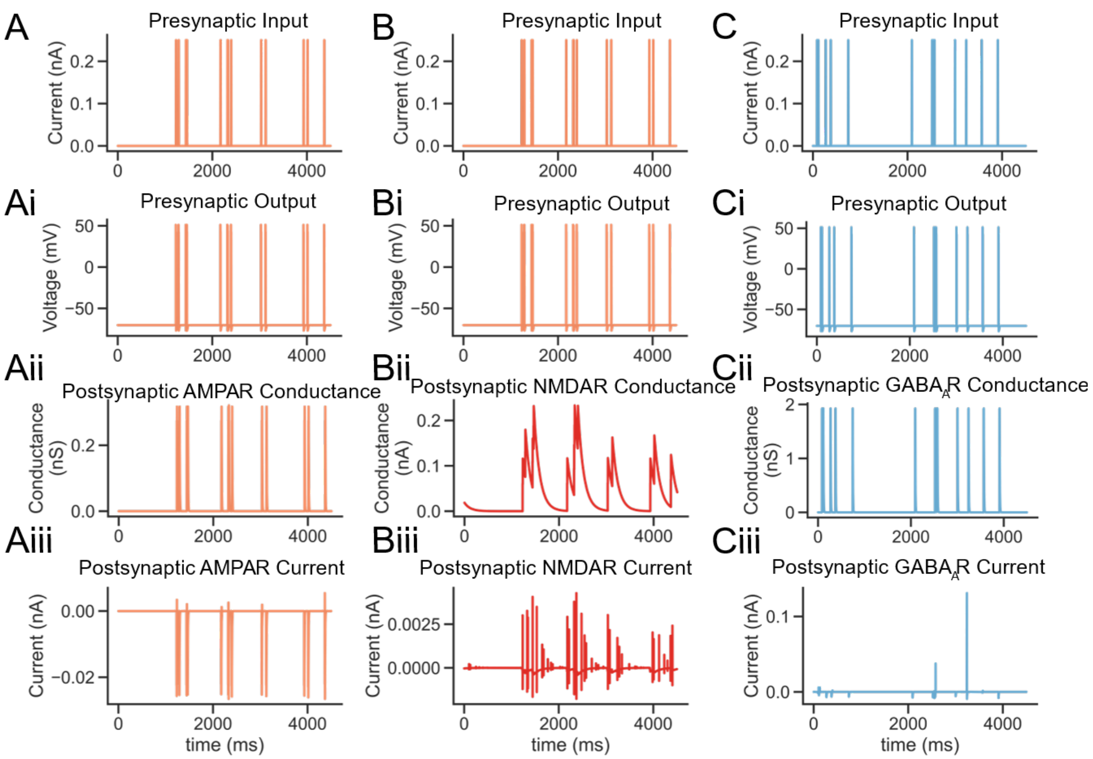


**Supplemental Figure 11. Presynaptic stimulation reliably evokes presynaptic spikes and postsynaptic currents in model dopamine neurons**. **A**. Stimulus delivered to presynaptic excitatory neurons produced high-fidelity presynaptic spikes (**Ai**) that produced postsynaptic AMPAR conductance (**Aii**) triggering postsynaptic AMPAR currents (**Aiii**). The same presynaptic excitatory neuron (**B, Bii**) also generated a postsynaptic NMDAR-mediated conductance and current (**Biii**). A separate set of presynaptic inhibitory neurons stimulated at a higher frequency (6 Hz Poisson, **C**) produced action potentials (**Ci**) that generated $GABA_AR$ conductance (**Cii**) producing $GABA_AR$ currents (**Ciii**).

**Supplemental Table 1. Channels and conductance values.**

| *Channel* | *Conductance(S/cm²)* | | |
|---|---|---|---|
| | *Somatic* | *Dendritic* | *Axonal* |
| *Na+ Markov 8 States* | 4 | 1.35 | 3 |
| *K+ Markov 5 States* | 4 | 4 | 6.5 |
| *Background Na+* | 0.000001 | 0.000001 | 0.000001 |
| *SK* | 0.005 | 0.005 | 0.005 |
| *A-type K+* | 0.0025 | 0.0175 | 0.0025 |
| *M-type K+* | 0.01 | 0.01 | 0.001 |
| *Leak K+* | 0.00001 | 0.00001 | 0.00001 |
| *L-type Ca2+* | 0.001 | 0.001 | 0.0001 |
| *N-type Ca2+* | 0.001 | 0.001 | 0.0001 |
| *T-type Ca2+* | 0.00075 | 0.00075 | 0.00075 |

**Supplemental Table 2. Western blot primary antibodies.**

| Primary Antibody Target | Producer, Product Number |
|---|---|
| Protein Kinase C-alpha (PKC α) | Cell Signaling Technologies (CST): #2056 |
| Phosphorylated PKC (pan; p-PKC βII Ser660 and homologs) | CST: #9371 |
| Phosphorylated (Ser) PKC Substrate | CST: #2261 |
